# Single-cell analysis of prenatal and postnatal human cortical development

**DOI:** 10.1101/2022.10.24.513555

**Authors:** Dmitry Velmeshev, Yonatan Perez, Zihan Yan, Jonathan E. Valencia, David R. Castaneda-Castellanos, Li Wang, Lucas Schirmer, Simone Mayer, Brittney Wick, Shaohui Wang, Tomasz Jan Nowakowski, Mercedes Paredes, Eric J Huang, Arnold R Kriegstein

**Author notes:** These authors contributed equally to this work (Dmitry Velmeshev, Yonatan Perez).

## Abstract

We analyze more than 700,000 single-nucleus RNA-seq profiles from 106 donors during prenatal and postnatal developmental stages and identify lineage-specific programs that underlie the development of specific subtypes of excitatory cortical neurons, interneurons, glial cell types and brain vasculature. By leveraging single-nucleus chromatin accessibility data, we delineate enhancer-gene regulatory networks and transcription factors that control commitment of specific cortical lineages. By intersecting our results with genetic risk factors for human brain diseases, we identify the cortical cell types and lineages most vulnerable to genetic insults of different brain disorders, especially autism. We find that lineage-specific gene expression programs upregulated in female cells are especially enriched for the genetic risk factors of autism. Our study captures the molecular progression of cortical lineages across human development.

**One Sentence Summary:** Single-cell transcriptomic atlas of human cortical development identifies lineage and sex-specific programs and their implication in brain disorders.

## Main text

Development of the human cerebral cortex spans months during prenatal stages and years after birth, generating tens to hundreds of cell types across multiple cortical areas. This complex process is orchestrated by lineage-specific gene expression programs that guide the production, migration, differentiation and maturation of neuronal and glial cell types, as well as the formation of projections and neuronal circuits. Alterations in these regulatory gene programs during development lead to the pathogenesis of neurodevelopmental and psychiatric disorders, including autism spectrum disorder (ASD) and schizophrenia (SCZ). Most previous studies have focused on investigating the molecular processes that underly human cortical development during the second trimester of gestation (*1–5*), the peak of cortical neurogenesis and neuronal migration. These studies have revealed molecular signatures of progenitor cells and neuronal and glial cell types, as well as the early specification of neurons into broad subtypes and their arealization across the cortex. However, later stages of human cortical development, including the third trimester of gestation, birth, and neonatal and early postnatal development, have been largely studied using bulk genomic approaches.

### Single-nucleus RNA sequencing analysis of prenatal and postnatal human cortical development

To gain a comprehensive view of human cortical development across prenatal and postnatal stages, we utilized single-nucleus RNA sequencing (snRNA-seq) (*6*) to profile 413,682 nuclei from 108 tissue samples derived from 60 neurotypical individuals. We sampled nuclei from ages spanning from the second trimester of gestation to adulthood, including samples from the third trimester and early postnatal stages that are often excluded or underrepresented in genomic studies of the human brain. We acquired data from the ganglionic eminences, the major source of cortical interneurons (*7, 8*), as well as from the cortex. We used Seurat (*9*) to perform unbiased clustering and UMAP embedding. After removing a cluster of cell debris (**Fig S1A**), we retained 358,663 nuclei. To extend our analyses to more brain samples and nuclei, we integrated our data with published datasets of prenatal and postnatal human cortical development (*10–12*). After data integration (**Fig S1B**), our final dataset included 709,372 nuclei and 169 brain tissue samples from 106 individuals (**Fig 1A**, **Table S1**). We identified clusters corresponding to neural progenitors, as well as the major subtypes of excitatory and inhibitory neurons, glia, and vascular cells (**Fig 1B-C**), indicating that we were able to capture transcriptomic changes underlying differentiation and maturation of cortical cell types across development. We detected similar numbers of genes, transcripts, and mitochondrial RNA ratios across different samples (**Fig S1C**) with a median of 1106 genes and 1609 transcripts per nucleus and some variability sample-to-sample. These relative numbers are comparable with published single-cell genomics data collected from the human brain (*13*), with mature neuron cell types expressing higher numbers of genes and transcripts than other cell types (**Fig S1D**). We did not observe batch effects, with nuclei from different samples well intermixed, and no clusters composed of nuclei from a single sample (**Fig S1E**). Nuclei were captured from the prefrontal, cingulate, temporal, insular and motor cortices (**Fig S1F**). For prenatal samples that were not sex-identified, we determined their sex using sex-specific gene expression (**Fig S1G**). Our dataset included 45 female and 61 male subjects. We observed that nuclei clustered according to developmental age (**Fig 1D**), suggesting that transcriptomic changes associated with development are a major driver of cell identity.

**Figure 1.**
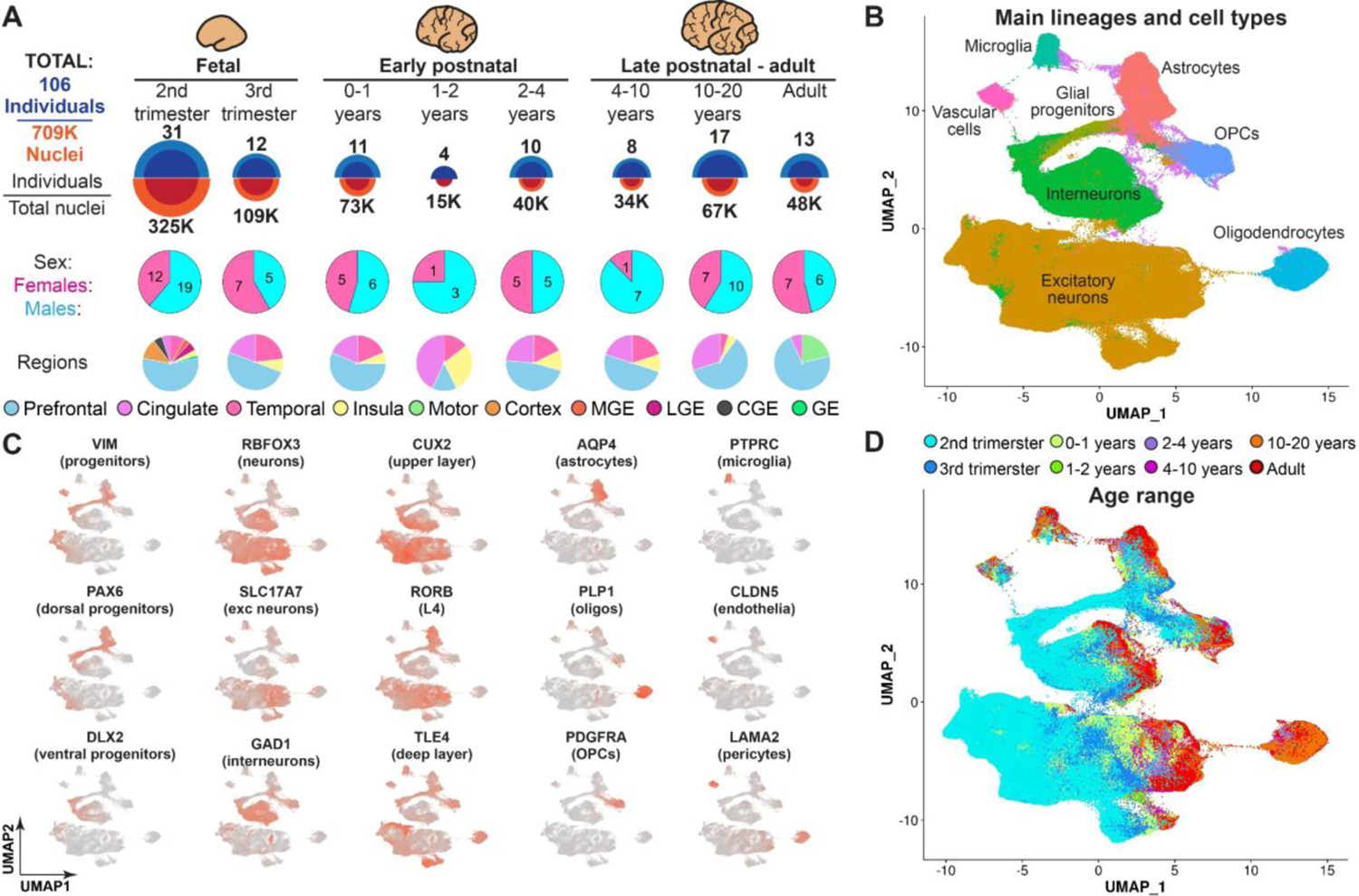
Brain tissue samples used for data collection and initial clustering of snRNA-seq data. **A)** Overview of the tissue samples used in the current study, including the number of individuals, as well as ages and brain regions captured in the snRNA-seq dataset. MGE-medial ganglionic eminence, LGE-lateral ganglionic eminence, CGE-caudal ganglionic eminence, GE-ganglionic eminence. **B)** Clustering of the entire dataset, with the major lineages labeled. **C)** Expression of cell type-specific markers used to determine cardinal lineages. **D)** Nuclei labeled by their developmental age.

### Analysis of specific excitatory neuron and interneuron lineages

We next examined the developmental trajectories of excitatory and inhibitory neurons. First, we selected clusters corresponding to dorsal forebrain progenitors (including radial glia and intermediate precursor cells) as well as clusters containing excitatory neurons. By re-clustering this data and referencing molecularly defined cell types annotated in the Allen Brain Atlas (*14*), we identified clusters corresponding to known subtypes of excitatory neurons, including upper (L2-3) and deep-layer intertelencephalic (L5-6-IT) projection neurons, layer 4 neurons (L4), layer 5 (L5) and layer 6 (L6) corticofugal projection neurons, as well as subplate neurons (SP) that were present transiently during the second trimester (**Fig 2A, Fig S2A**). We next used monocle 3 (*15*), as well as custom scripts (see **Methods**) to construct cellular trajectories based on snRNA-seq data (**Fig 2A; Fig S2B**), select trajectory branches corresponding to specific lineages, and calculate pseudotime for each nucleus. Pseudotime corresponded well to the developmental age of nuclei in each lineage (**Fig 2A**). We identified several branching points in the trajectory: between two major groups of excitatory neurons: L2-3, L4 and L5-6-IT (Ex1) and L5 and L6 (Ex2), as well as between L4 and L2-3/L5-6-IT (Ex3). Next, we aimed to investigate developmental gene expression changes during differentiation and maturation of GABAergic interneuron (IN) lineages. We selected nuclei from ventral forebrain progenitors, as well as cortical interneurons, re-clustered the data and identified known classes of cortical interneurons (**Fig 2B, Fig S2C**), including interneurons expressing VIP, calretinin (CALB2), reelin (RELN), and nitric acid synthase (NOS), and chandelier (PV-CH) and basket (PV-BSK) interneurons expressing parvalbumin, *MME* and *TAC1*, as well as interneurons expressing somatostatin (SST) and co-expressing *SST* and reelin (SST-RELN). We then reconstructed lineage trajectories corresponding to each interneuron subtype (**Fig 2B, Fig S2D**), as well as point of trajectory divergence, such as trajectory branches including MGE-(IN1) and CGE-derived (IN2) interneurons. We calculated pseudotime for each nucleus, which correlated well with the developmental age of the interneurons. Next, we asked whether different neuronal lineages in the human cortex mature at different rates. We correlated pseudotime with the developmental age in each neuronal lineage and observed that neuronal types fell into two main groups: those that mostly matured by the end of the second trimester, and those whose transcriptome profiles continued to change through the third trimester and after birth (**Fig 2C**). The first group included L5, L5-6-IT and all interneuron subtypes, whereas the second group contained L2-3, L4 and L6 excitatory neurons. This result suggests that certain types of human cortical neurons have a protracted maturation timeline.

**Figure 2.**
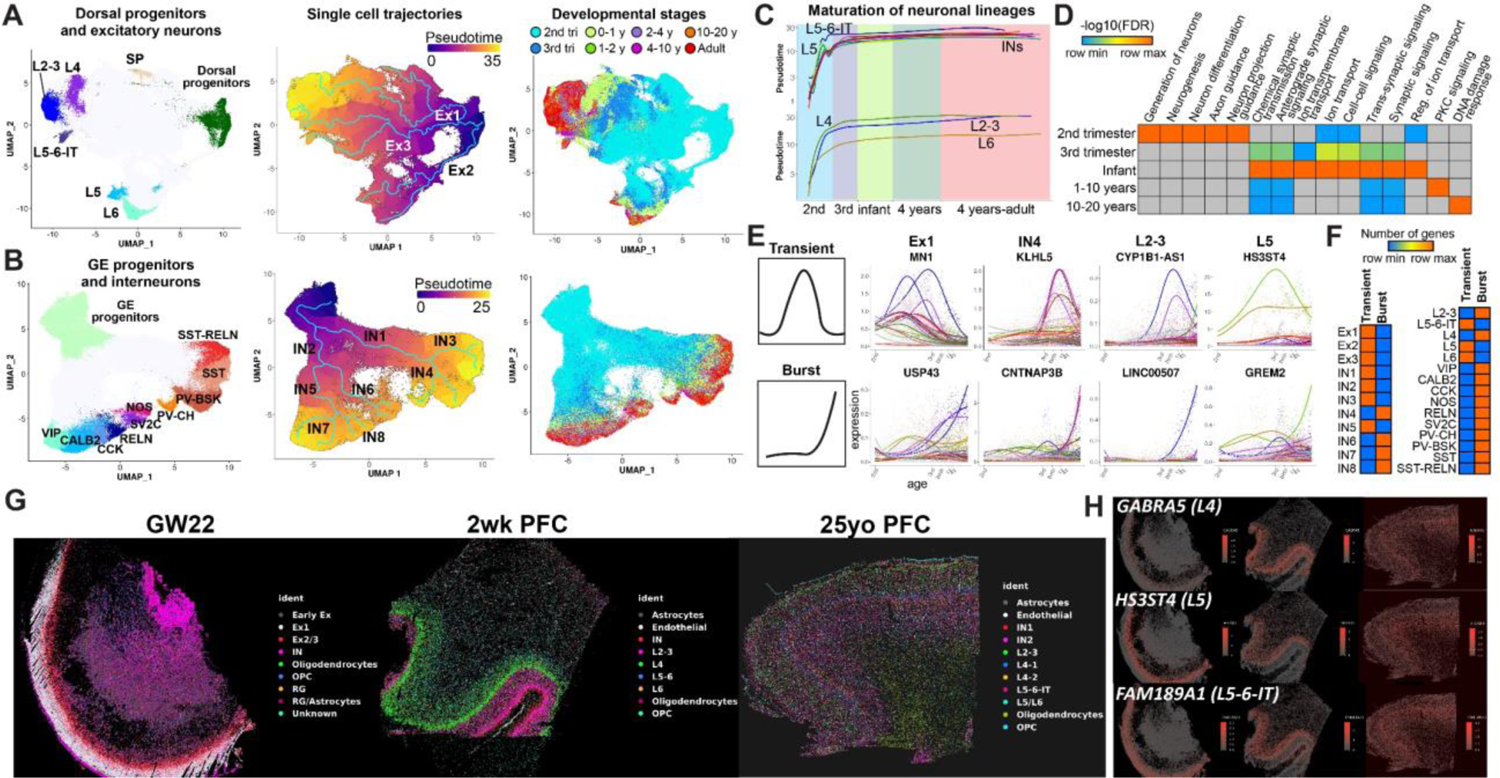
Analysis of excitatory and inhibitory neuron lineages. **A)** Cell types, reconstructed single-cell trajectories, and age distribution for subtypes of excitatory neurons. L2-3 – upper-layer cortico-cortical projection neurons, L4 – layer 4 neurons, L5-6-IT – deep-layer intratelencephalic projection neurons, L6 – layer 6 neurons, L5 – layer 5 neurons, SP – subplate neurons. **bB**Identification of interneuron trajectories. **C)** Rates of maturation of subtypes of excitatory neurons and interneurons. **D)** Gene ontology analysis of genes with different age of onset of expression. **E)** Examples of top lineage and branch-specific genes with transient and burst expression patterns. **F)** Number of transient and burst genes in specific lineages and branches. **G)** Spatial transcriptomic analysis of 140 lineage-specific genes, showing the spatial map of annotated cell-types across development (GW22 = 22 weeks of gestation; 2wk = 2 weeks postnatal; 25yo = 25-year-old; PFC = prefrontal cortex). **H)** Examples of deep-layer neuronal markers with early patterned layer-specific expression (putative layer location is in brackets).

Once we isolated trajectory branches corresponding to each neuronal lineage, we sought to identify lineage-specific gene expression programs. We employed an approach that allows identification of lineage-specific programs by comparing dynamic expression profiles of each gene in a lineage of interest to all other neuronal, glial and non-neural lineages ( see **Methods**). In addition, we applied this approach to identify genes specific to related lineages in the excitatory neuron and interneuron trajectory branches. In total, we identified 1062 lineage-specific genes and 405 branch-specific genes (**Table S2**). We classified these genes based on the age of onset of gene enrichment (50% of the maximum expression) and performed gene ontology analysis for the genes upregulated at each developmental timepoint (**Fig. 2D**). During the second trimester of gestation, we saw enrichment in pathways related to neurogenesis, differentiation, and process growth. Upregulation of synaptogenesis and ion transport pathways could be observed during the third trimester but was most profound between birth and one year of age. Enrichment in synaptic pathways could be observed until adulthood.

In addition to classifying genes based on their age of appearance, we also characterized dynamic expression patterns of lineage-specific genes. The two most common patterns we observed were transient expression and burst expression where upregulation would start at a certain age and continue into adulthood (**Fig 2E**). Our analysis identified several putative regulators of neuronal lineage commitment, such as transcriptional regulator *MN1* specific to L2-3, L5-6-IT and L4 neurons, noncoding RNAs *CYP1B1-AS1* and *LINC00507* enriched in L2-3 neurons, and *HS3ST4* specific to L5 neurons. We saw that genes enriched in more broad lineage branches tended to be transiently expressed genes, whereas genes specific to mature neuronal cell types mostly followed burst expression patterns (**Fig 2F**). This suggests gradual commitment and specification of neuronal cell types through a series of transient and burst transcriptional events. We also classified additional less common expression patterns, such as biphasic expression (**Fig S2e**) and identified different biological processes enriched for genes with burst and transient expression patterns (**Fig S2F**). Finally, we identified genes dynamically expressed during the specification of subplate neurons by comparing lineages during the second and third trimester of gestation when these cells are present (**Fig S2G**). Using spatial transcriptomic analysis of 140 genes across three developmental timepoints, we were able to identify and visualize the spatial location of cell-specific clusters overlaid on the tissue cytoarchitecture. Focusing on early-emerging lineage-specific genes, we validated the spatiotemporal expression of excitatory layer-specific markers **(Fig 2G-H, Fig S3)**. We observed that broad classes of excitatory neurons in the Ex1, Ex2 and Ex3 trajectory branches are restricted to specific cortical layers during the second trimester of gestation. Moreover, several markers of L4 neurons, such as *HPCA* and *GREM2*, are expressed in a layer-restricted manner during the second trimester of gestation suggesting that L4 neuronal identity starts to be specified early in development. The layer identity of most excitatory neurons emerges by birth (**Fig S3**) based on the lineage-specific signatures that we find specify human cortical neurons and their segregation to cortical layers.

### Dissection of glial and non-neural lineages

We further focused on the analysis of glial lineages, including astrocytes and oligodendrocytes. We re-clustered glial progenitors, oligodendrocyte precursor cells (OPCs), oligodendrocytes, and astrocytes and performed trajectory analysis (**Fig 3A**). We identified two types of astrocytes: fibrous astrocytes with high expression of *GFAP*, and protoplasmic astrocytes with low expression of *GFAP* and high expression of glutamate transporter GLAST (*SLA1A3*) (**Fig S4A**). Next, we performed identification of lineage-specific genes in the manner described for neuronal lineages (**Table S2**). We first focused on genes that were expressed at the divergence of astrocyte and oligo trajectory branches (**Fig 3B**). We observed well-known transcription factors guiding commitment to the oligo and astrocyte lineages, including *OLIG1*, *OLIG2*, *ID4* and *SOX9*, as well as other putative regulators, such as the zinc finger protein ZCCHC24 specific to the oligo lineage and a DNA binding protein STOX1 enriched in astrocytes. When comparing fibrous and protoplasmic astrocytes, we identified gene programs specific to these cell types (**Fig 3C**). Genes upregulated in protoplasmic astrocytes after birth and during the first year of life were mostly associated with the transport of glutamate and its metabolites, suggesting a maturation program to support neuronal firing during the early postnatal period. For oligodendrocytes we observed that genes upregulated during the second and third trimesters were associated with glial cell differentiation, whereas myelination genes were upregulated after birth and continued to be expressed into adulthood (**Fig 3E**). Analysis of microglia development (**Fig 3F**) identified three cell trajectories (MG-1-3), one of which (MG-3) was associated with highly activated microglia and was present in a small number of samples. These trajectories were confirmed by an alternative analysis using Slingshot (**Fig S4B**) (*16*). We focused on the non-activated microglia trajectories (MG-1 and MG-2) which were differentiated from each other by expression of a pro-inflammatory microglia marker, *IKZF1,* expressed in MG-2. *IKZF1* was the only gene differentiating MG-1 and MG-2, suggesting that these trajectories may represent two different states of the same microglia cell type rather than different subtypes; therefore, we focused on genes developmentally expressed in both of these microglia cell clusters. By performing Gene Ontology (GO) analysis of microglia-specific genes upregulated at different developmental stages, we observed complement genes associated with synaptic pruning upregulated in microglia after birth and during the first year of life (**Fig S4C**, **Fig 3G**). These findings suggest that the developmental period between birth and one year of life is a critical period of synaptic formation and plasticity that involves not only neuronal lineages, but also protoplasmic astrocytes and microglia. Finally, we identified gene programs associated with the maturation of brain endothelial cells and pericytes (**Fig S4D-F**). Our data suggests a coordinated maturation of neuronal and glial cell functions that insures proper formation and maintenance of neuronal circuits.

**Figure 3.**
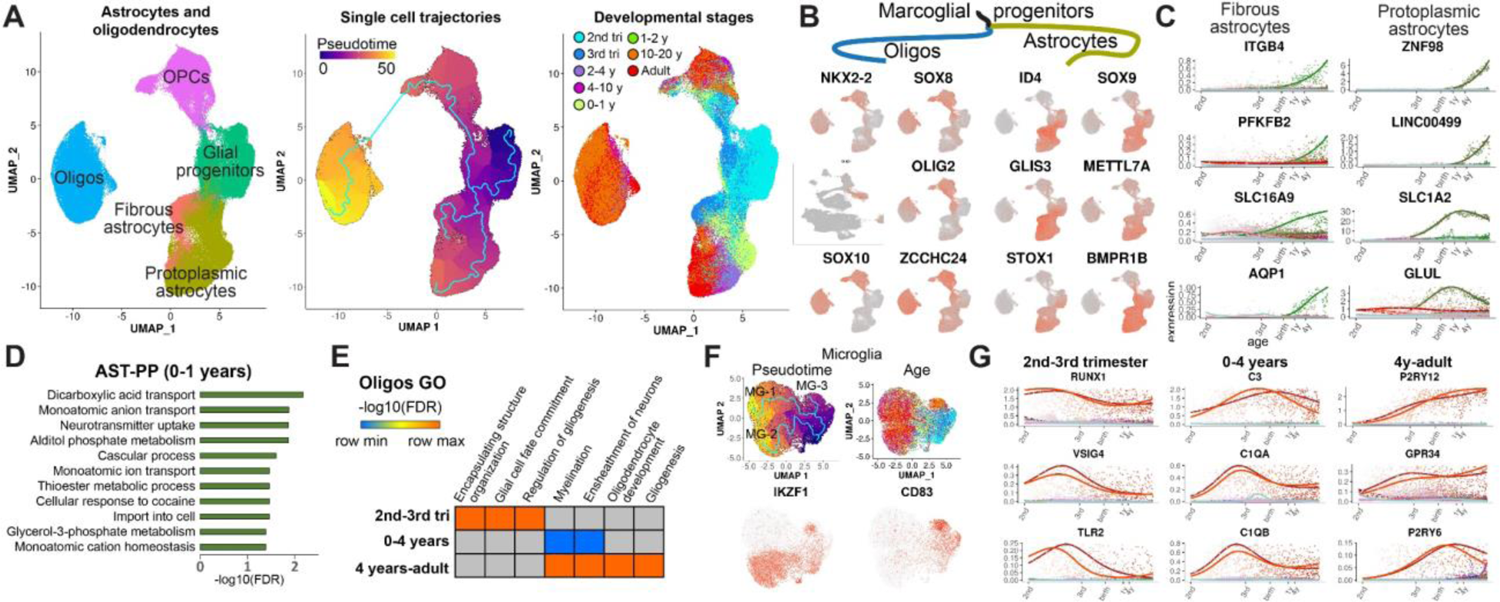
Analysis of inhibitory cortical interneuron lineages. **A)** Clusters and trajectories of glial progenitors, astrocytes and oligodendrocytes. **B)** Sample genes specific to oligodendrocyte and astrocyte lineage branches. **C)** Examples of top dynamically expressed genes specific to fibrous and protoplasmic astrocytes. **D)** Gene ontology analysis of protoplasmic astrocyte-specific genes expressed during the first year of life. **E)** Pathways enriched for oligo lineage-specific genes expressed at different developmental stages. **F)** Analysis of microglia lineages. **G)** Temporal patterns of developmental microglia genes.

### Integration of with single-cell open chromatin data and identification of lineage-specific gene regulatory networks

Epigenetic regulation plays a crucial role in cortical neuron lineage commitment and specification. In order to identify lineage-specific transcriptional and epigenetic regulators of the cortical lineages identified in the snRNA-seq data, we leveraged recently published single-nucleus ATAC-seq (snATAC-seq) data from the developing human cortex during prenatal and postnatal stages (*10, 11, 17, 18*). First, we combined snATAC-seq data from four datasets, obtaining 290,239 snATAC-seq profiles from 57 tissue sample and 42 individuals across the second trimester, early postnatal stages of development, as well as adult life. We then utilized Seurat to integrate the resulting snATAC-seq data with our snRNA-seq data and mapped the integrated snATAC-seq data to the snRNA-seq clusters, UMAP space, and cell types (**Fig 4A, see Methods**). We observed that the developmental ages for the snATAC-seq and snRNA-seq profiles are well aligned (**Fig 4A**, **Fig 1D**). Gene activity (open chromatin in the promoter and gene body) of cell type marker genes suggested that snATAC-seq profiles mapped to corresponding transcriptionally defined neuronal and glial cell types (**Fig S5A**). Next, we repeated the integration and mapping procedure for three major lineage classes: excitatory neurons, interneurons, and glia (astrocytes and oligodendrocytes) (**Fig 4B-D, Fig S5B-D**). We omitted microglia and vascular cells due to a low number of snATAC-seq profiles in these lineages. After mapping snATAC-seq data to the transcriptionally defined lineages, we selected snATAC-seq cells along each lineage branch (**Fig S5B-D)**. Not all lineages could be reliably recovered due to the smaller size of the snATAC-seq dataset and the lack of key developmental stages, such as the third trimester. We therefore focused on lineages that had ATAC cells along the entire span of the trajectory, including four excitatory neuron lineages, five interneuron lineages, and both types of astrocytes and oligodendrocytes as indicated in **Fig 4B-D**. Plots of lineage-specific gene activity over pseudotime demonstrated that we accurately mapped and selected lineage-specific snATAC-seq profiles (**Fig 4B-D)**. Finally, we leveraged SCENIC+ (*19*), a recently developed algorithm that uses paired single-cell transcriptomic and open chromatin data to identify enhancer gene regulatory networks (eGRN) and candidate transcription factors that regulate expression of target genes in these networks. We applied SCENIC+ to the snRNA-seq and snATAC-seq profiles in each lineage to identify open chromatin regions correlated with pseudotime, putative enhancers, candidate transcription factors (TF) that bind them, and their association with lineage-specific dynamically expressed genes (**Table S3**). In total, we identified 42 transcription factors regulating 1373 lineage-specific genes through predicted binding of 4846 regulatory chromatin regions. We observed networks regulated by previously known lineage-specific transcriptional regulators, such as SOX5 in deep-layer projection neurons (**Fig 4B**), LHX6 in MGE-derived PV and SST interneurons (**Fig 4C, Table S3**), OLIG2 in oligodendrocytes and SOX9 in astrocytes (**Fig 4D**). Additionally, we identified previously unrecognized (at the best of our knowledge) putative lineage-specific transcriptional regulators, such as BACH2, predicted to regulate several key deep-layer transcription factors in L5 neurons, including FOXP2 and FEZF2, as well as NFIX and ZNF184 specific to L2-3 neurons and regulating expression of the upper-layer master transcription factor, CUX2 (**Fig 4B**). Our results also suggest the role of the transcription factor MAFB in parvalbumin interneuron specification (**Fig 4C**), as well as of FOXN2 and RFX4 in determining the fate of oligodendrocytes and protoplasmic astrocytes, respectively (**Fig 4D**). Our data sheds new light on epigenetic control of neural lineage commitment and identifies putative transcription factors and regulatory networks that define the fate of specific human cortical neuronal and glial cell types.

**Figure 4.**
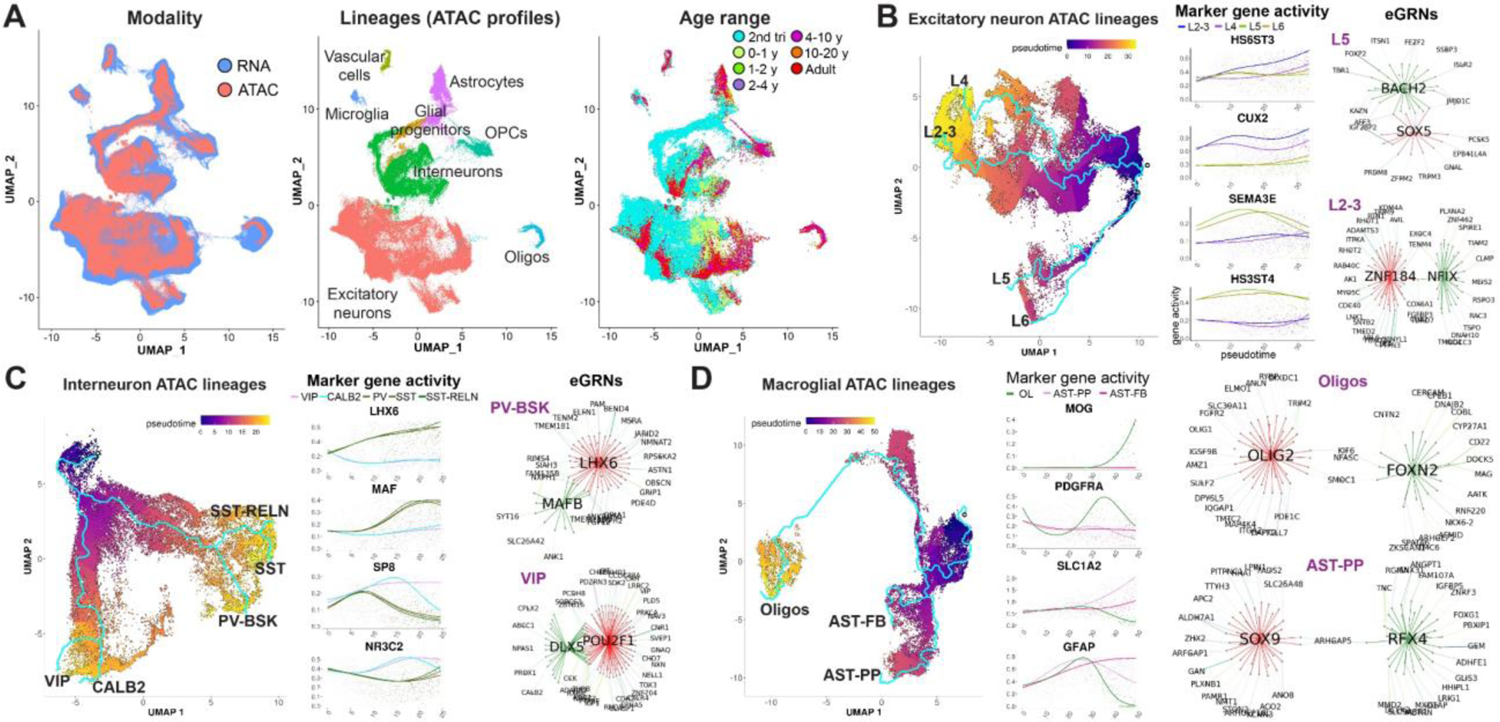
Identification of lineage-specific epigenetic and transcriptional regulators. **A)** Integration of snRNA-seq and scATAC-seq data. scATAC-seq data was mapped on the scRNA-seq coordinates, clusters and cell types. **B-D)** Analysis of enhancer gene regulatory networks (eGRNs) in excitatory neuron lineages **(B)**, as well as interneurons **(C)** and glial lineages **(D)**. Network plots (eGNRs) display transcription factors predicted to bind enhancer regions to regulate lineage-specific transcriptional programs. Edge colors indicate regulation by different transcription factors. Top 20 genes based on the predicted confidence of interaction are shown for each transcription factors network.

### Identification of region and sex-enriched lineage-specific gene programs

Since we sampled our transcriptomic data from different cortical regions, we asked whether lineage-specific developmental gene expression profiles might be spatially defined, and vary depending on cortical area. We focused on the frontal/prefrontal cortex (PFC) since we had the most complete sampling of this cortical area across developmental stages (**Fig S6A**). We compared each neuronal and glial lineage trajectory in the PFC to the trajectories in all other cortical areas and identified PFC-enriched developmentally regulated genes in each lineage (**Table S4**). We observed more PFC-specific genes in excitatory neuron lineages, especially in intertelencephalic upper (L2-3) and deep-layer (L5-6-IT) neurons, as well as in astrocytes and oligodendrocytes, whereas most interneuron lineages and microglia expressed fewer PFC-specific genes (**Fig S6B**). After performing GO analysis for PFC genes specific to neuronal lineages, we observed enrichment in cell adhesion and synaptic transmission pathways (**Fig S6C**). Analysis of glia-specific PFC genes demonstrated enrichment in different categories of biological pathways associated with cell division and cell migration (**Fig S6D**). Examples of neuronal PFC genes included synaptojanin 2 binding protein *SYNJ2BP* regulating receptor localization and signal transduction at the synapse and the cation channel *TRPC7* (**Fig S6E**). PFC fibrous astrocytes upregulated R-spondin 2 (*RSPO2*) and Frizzled Class Receptor 8 (*FZD8*), which both participate in Wnt signaling and cell migration. Our results suggest cortical areal differences in lineage-specific transcriptomic programs, with synaptic genes upregulated in neuronal cell types and cell division and cell migration programs activated in glial cells in the developing PFC. PFC-specific expression of synaptic genes in neuronal cell types suggests regional specification of neuronal circuits during development.

We next asked whether the development of specific cellular lineages is modulated in a sex-dependent manner. For each lineage analyzed, we isolated female and male nuclei (**Fig 5A**, **Fig S7A**) and identified dynamically expressed genes enriched during either female or male development. In total, we identified 740 female-enriched genes and 312 male-enriched genes (**Table S5**). Only a small fraction of male genes showed female/male enrichment in a lineage-specific manner (20/312, 6.4%), whereas more than half of female genes showed lineage specificity of sex enrichment (510/740, 69%). Despite several top female-enriched genes located on X and Y chromosomes (including *XIST* and *PCDH11Y*), sex-enriched genes were evenly distributed across all chromosomes (**Fig S7B**), suggesting that sex-dependent developmental modulation of gene expression is not directly dependent on transcription from the sex chromosomes. We next performed GO analysis of female and male-enriched genes, focusing on the neuronal, astrocyte and oligodendrocyte lineages where we had large number of samples and nuclei from both sexes. We observed substantial difference between the biological processes associated with female and male-enriched genes: female genes were involved in developmental processes, including cell adhesion, CNS development, synaptic transmission and membrane potential regulation (**Fig. 5B**), whereas male genes were associated with RNA metabolism and translation (**Fig. 5C**). Only a small number of male-specific genes such as *YBX1* and *LINGO1* were associated with developmental processes; however, these genes were enriched across multiple male lineages (**Fig S7C**). We classified sex-enriched genes according to their dynamic expression pattern and saw that the majority were expressed transiently (**Fig. 5D)**, with over 90% having peak expression during the second trimester (**Table S5**). This suggest early and transient sex-dependent developmental modulation of cortical lineages. Sex-enriched genes were more abundant in excitatory neuron lineages compared to interneurons (**Fig 5E**) and were also abundant in female fibrous astrocytes. Several top lineage-specific female-enriched genes were associated with neuronal, glial and endothelial development (**Fig 5F, Fig S7D**). These included nuclear hormone receptor/transcription factor *RORA* in L2-3 neurons, synaptic protein neurexophilin 3 (*NXPH3*) in L6 neurons, transcription factor *HES4* in fibrous astrocytes, and an actin filament depolymerization enzyme, *MICAL3,* in oligodendrocytes. Overall, our results point to modulation of neuronal and glial developmental programs during second trimester female brain development.

**Figure 5.**
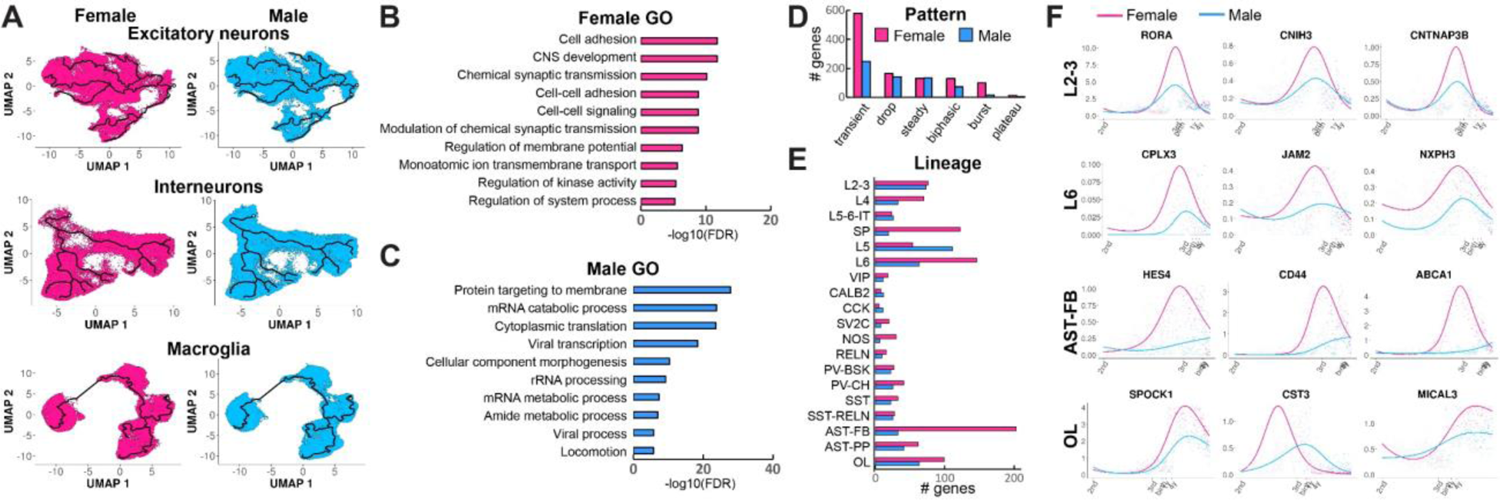
Analysis of sex-specific developmental programs in human cortex. **A)** Female and male developmental trajectories of excitatory neurons, interneurons, astrocytes and oligodendrocytes. **B-C)** Gene ontology analysis of female and male-enriched genes. **D)** Dynamic expression patterns of sex-enriched genes. **E)** Sex enrichment of developmental gene expression across neuronal and glial lineages. **F)** Examples of top female-enriched genes in specific lineages.

### Enrichment of lineage-specific developmental gene programs for risk factors of brain disorders

Once we defined lineage and sex-specific developmental gene programs in human cortical cell types, we sought to investigate how these transcriptional programs may be affected in neurodevelopmental, psychiatric, and neurodegenerative disorders. We compiled all lineage-specific gene signatures for excitatory neurons, astrocytes, oligodendrocytes, interneurons, microglia, endothelial cells and pericytes, in total obtaining 2796 unique genes, and divided them into 5 groups based on their age of expression onset (50% of max expression). We then overlapped this gene list with lists of rare gene variants associated with the risk of ASD from the Simons Foundation Autism Research Initiative (SFARI) Gene database (*20*), as well as GWAS genes for the risk of SCZ (*21*), bipolar disorder (BPD) (*22*) and Alzheimer’s disease (AD) (*23*) (**Fig 6A, Table S6**). We observed a large enrichment for genes associated with risk for ASD, SCZ and BPD in the second trimester, with expression of ASD and BPD risk genes extending to the third trimester. The risk of neurodevelopmental disorders dropped during later stages of development. Expression of ASD risk genes remained mostly flat and only slightly above the significance level, demonstrating a pattern different from neurodevelopmental and psychiatric disorders. We next analyzed enrichment of disease risk genes across cortical lineages (**Fig 6B**). We were able to detect significant enrichment for ASD risk genes in L5-6-IT and L5 neurons, whereas AD risk genes were enriched in microglia. We focused on ASD since we observed the strongest enrichment for the risk of this disorder among developmentally regulated genes, and because a large amount of genetic risk data is available for this disorder. We observed developmental enrichment of ASD risk genes with SFARI score 2 and 3 but not score 1 and did not find enrichment in syndromic ASD genes (**Fig 6C**). We observed a significant enrichment among high-confidence ASD risk genes (ASD-HC) based on the TADA analysis (*24*). We conclude that the genetic burden of ASD has the potential to affect the development of specific neuronal cell types, especially deep-layer intertelencephalic projection neurons and L5 neurons. We next explored enrichment of ASD risk genes in sex-specific developmental programs. We observed strong enrichment of female-specific developmental genes in both SFARI and HC-ASD gene lists (**Fig 6D**). Male-specific genes were less frequently found among SFARI genes, and we did not find a meaningful overlap between male-enriched and high-confidence ASD genes. This finding points to a strong enrichment of the genetic risk of ASD among developmental genes that are more highly expressed in female cells. SFARI genes were enriched in female cells across multiple neuronal cell types, especially the subplate and L6 excitatory neurons, as well as oligodendrocytes and fibrous astrocytes, but not in microglia or vascular cell types (**Fig 6E**). This suggests a role of the subplate in the pathogenesis of ASD. Examples of female-specific high-confidence ASD risks genes included the subplate-specific transcription factor *NR4A2* and the neuronal transcription factor *MEF2C* that were upregulated in female subplate cells, as well as a regulator of axon guidance and synaptogenesis, neurexin 2 (*NRXN2),* and *PCDH15* encoding a cell adhesion molecule in female L6 neurons (**Fig 6G**). Our findings provide strong evidence supporting the ASD female protective effect hypothesis (*25*), and suggest that fine-tuning of cortical cell lineages by sex-specific developmental programs can contribute to the male bias in the pathogenesis of ASD.

**Fig 6.**
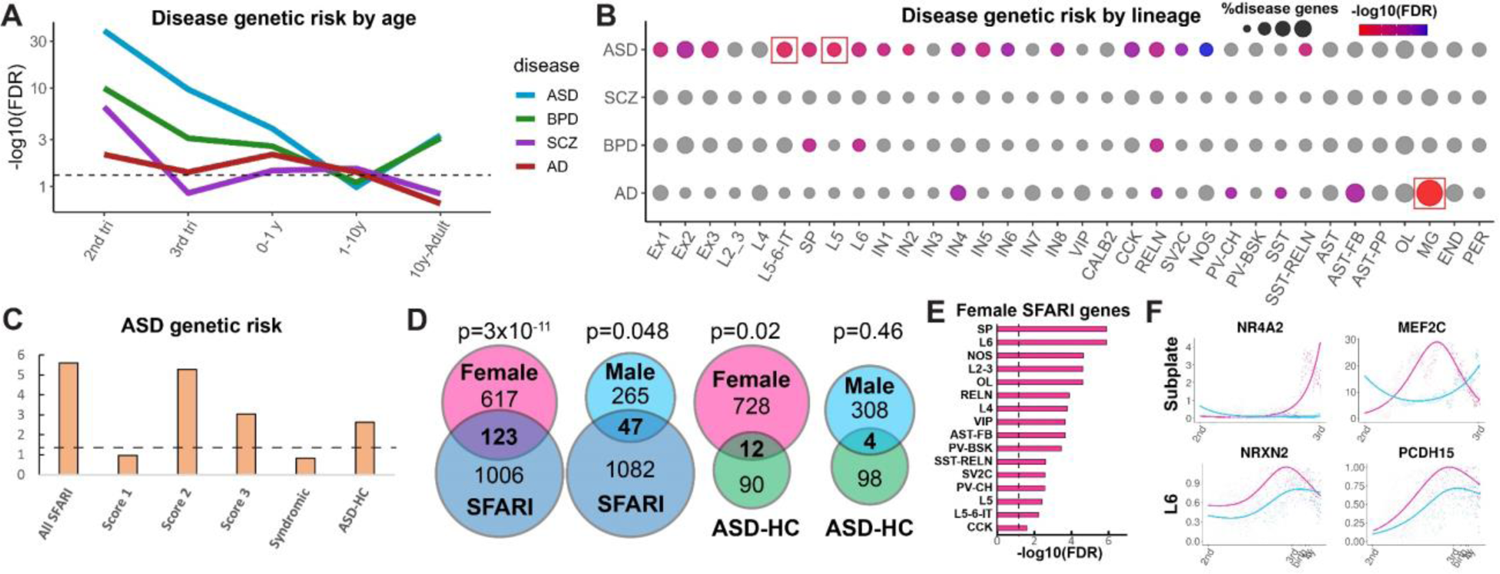
Lineage enrichment of ASD risk genes. **A)** Enrichment of disease risk genes across developmental stages. **B)** Disease risk gene enrichment across lineages and lineage branches of neuronal, glial, and vascular cell types. Red squares indicate statistical significance. **C)** Enrichment of lineage-specific developmentally regulated ASD risk genes of different categories and evidence scores. **D)** Overlap between ASD risk genes and female and male-enriched developmental gene programs. **E)** Enrichment of sex-specific genes across specific lineages. **F)** Temporal patterns of female-enriched genes that are known risk factors for ASD.

## Discussion

By generating single-nucleus RNA-seq data from the developing human cortex and integrating the findings with previously published datasets, we performed a large-scale unbiased transcriptomic analysis of human cortical development throughout the lifespan. By reconstructing single-cell trajectories and identifying genes that are expressed in a lineage-specific manner we created a compendium of developmental programs for all the major cortical cell types. By integrating our data with published single-cell chromatin accessibility datasets, we identified enhancer-gene regulatory networks and transcription factors that are predicted to control the commitment and differentiation of specific cortical neural lineages. In addition, we characterized sex and brain region-specific gene programs that are used by specific lineages of cortical cell types. We find that female-enriched genes are associated with neurodevelopmental processes, whereas male-enriched genes are involved in protein translation control, suggesting sex-specific variation of developmental trajectories. We also find that developmental gene programs utilized by cortical excitatory neurons, astrocytes and oligodendrocytes are the most region-specific. Interneurons, in contrast, express few region-specific genes during development, consistent with data on regional signatures of cortical cell types in the mature human brain (*26*).

We investigated the enrichment of genetic risk factors for brain disorders, focusing on ASD, and found that the developmental programs of both deep-layer intratelencephalic and corticofugal projection neurons are enriched for ASD risk genes. These data are in agreement with previous reports of enrichment of ASD genes in deep-layer cortical neurons during mid-gestation (*27, 28*) but also suggest that both deep-layer neurons projecting to other cortical areas and to subcortical locations could be affected. We previously reported that upper-layer cortical excitatory neurons are most dysregulated in the cortex of idiopathic ASD patients (*29*). It would be an important future direction to elucidate how changes in pan-excitatory neuron programs during development can culminate in dysfunction of specific cortical neuronal populations, such as L2-3 neurons. It would also be valuable to explore whether the molecular pathology of upper-layer neurons is specific to idiopathic ASD, and whether it is driven by common gene variants, rather than rare variants with strong effect sizes (*30*). In addition, we observed a strong enrichment of ASD genetic risk factors among female-specific developmental genes. Since these female-enriched ASD risk genes have higher expression in females during cortical development, is possible that this higher baseline expression renders female brain more resistant to genetic insults causing autism, especially to haploinsufficiency that can reduce transcript or protein expression by affecting one of the two alleles. This finding might explain the 4:1 male to female ratio of individuals affected by ASD and suggests the importance of sexual dimorphism in human brain development. However, the role of sex hormones in the increased male to female ratio in ASD is not to be discounted, and additional studies are needed to reconcile the role of early development and later sex-specific processes in the pathogenesis of autism. Our preliminary findings indicate the cell type-specific risk of BPD and SCZ, but more detailed genetic studies are needed to further dissect cell type and developmental stage vulnerability. The data generated here may help enable fine-grained understanding of human brain development and provide insight into mechanisms of neurodevelopmental disorders.

Our study, however, is limited by the technical difficulty of integrating snRNA-seq and scATAC-seq data as well as by the lack of inclusion of earlier developmental stages, such as the first trimester, due to challenges of integrating scRNA-seq and snRNA-seq datasets. Overcoming these obstacles will allow for even more comprehensive future understanding of how specific human cortical lineages develop. Moreover, single-cell epigenetic analyses of human brain development would be necessary to determine whether imprinting plays a role in regulating sex enrichment of developmentally expressed genes.

## Materials and methods summary

Brain tissue samples were sectioned using a cryostat to collect coronal cortical sections, lysed and ultracentrifuged to isolate nuclei. Nuclei were captured using 10x Genomics Single Cell 3’ v.2 kits.

Raw sequencing data were processed using 10x Genomics CellRanger and aligning reads to unsliced human transcriptome to capture reads from premRNAs. Dataset integration was performed using Harmony based on 10x chemistry, and clustering and UMAP embedding was carried out with Seurat. Monocle 3 was used to reconstruct lineage trajectories, and custom scripts were used to identify lineage-specific dynamically expressed genes (Supplementary Materials).

scATAC-seq data were integrated with snRNA-seq data using canonical correlation analysis in Seurat, after which different scATAC-seq chemistries were integrated using Harmony. Enhancer gene regulatory networks were identified using SCENIC+.

## Acknowledgments

We thank the NIH’s Brain Research through Advancing Innovative Neurotechnologies Initiative - Cell Census Network (BICCN) and The Brain Cell Data Center (BCDC) at the Allen Institute for Brain Science, as well as all of its members for the support of this work and helpful discussions.

## Authors contributions

DV designed the project, acquired tissue collection, performed nuclei isolation and 10x Genomics capture, analyzed the data and wrote the manuscript. YP performed spatial transcriptomic experiments, nuclei isolation and 10x Genomics capture and edited the manuscript. ZY helped with the analysis code and performed analysis of the oligodendrocyte lineage. JEV and DRC performed probe design and analysis of spatial transcriptomics data. LW performed spatial transcriptomic experiments. LS acquired adult samples, performed nuclei isolation and 10x Genomics capture. SM performed nuclei isolation and 10x Genomics capture. BW developed Cell Browser visualization tools. SW acquired second trimester tissue samples. TJN performed regional tissue dissections. MP and EJH acquired third trimester and early postnatal samples. ARK designed and supervised the project and edited the manuscript. All authors read the manuscript.

## Data and materials availability

Raw data can be accessed at the NeMo Archive, accession number nemo:dat-3ah9h9x (https://assets.nemoarchive.org/dat-3ah9h9x). Analyzed data (cell-count matrix and metadata) can be accessed through the UCSC Cell Browser, collection human-cortical-dev (https://pre-postnatal-cortex.cells.ucsc.edu), and at cellxgene (https://cellxgene.cziscience.com/collections/bacccb91-066d-4453-b70e-59de0b4598cd). Code is available at https://github.com/velmeshevlab/dev_hum_cortex and https://doi.org/10.5281/zenodo.7245297.

## Supplementary Materials

### Materials and Methods

#### Sample acquisition and selection

Samples were acquired from three different sources. 1) De-identified second-trimester tissue samples were collected at the Zuckerberg San Francisco General Hospital with previous patient consent in strict observance of the legal and institutional ethical regulations. Protocols were approved by the Human Gamete, Embryo, and Stem Cell Research Committee (institutional review board) at the University of California, San Francisco. These fresh tissue samples were dissected and snap-frozen in isopentane on dry ice. 2) De-identified second-trimester, third trimester and early postnatal tissue samples were obtained at the UCSF Pediatric Neuropathology Research Laboratory led by Dr. Eric Huang. These samples were acquired with patient consent in strict observance of the legal and institutional ethical regulations and in accordance to research protocols approved by the UCSF IRB committee. These samples were dissected and snap-frozen either on a cold plate placed on a slab of dry ice or in isopentane on dry ice. 3) Banked de-identified second-trimester, third trimester, early postnatal and adult tissue samples were obtained from the University of Maryland Brain and Tissue Bank through the NIH NeuroBioBank.

For postnatal ages, samples from individuals with known history of brain disorders or brain trauma were excluded from downstream analyses. For prenatal samples, samples with unusual neuropathology following pathological examination, as well as samples positive for commonly tested chromosomal aberrations, were excluded. Prior to performing nuclei isolation and single-nucleus RNA sequencing, samples were screened for RNA quality by collecting 100um-thick cryosections, isolating total RNA and measuring RNA Integrity Number (RIN) using the Agilent 2100 Bioanalyzer instrument. Only samples with RIN >= 6.5 were included in the study.

#### Nuclei isolation and generation of single-nucleus RNA-seq data using 10x Genomics platform

40 mg of sectioned brain tissue was homogenized in 5 mL of RNAase-free lysis buffer (0.32M sucrose, 5 mM CaCl^2^, 3 mM MgAc^2^, 0.1 mM EDTA, 10 mM Tris-HCl, 1 mM DTT, 0.1% Triton X-100 in DEPC-treated water) using glass dounce homogenizer (Thomas Scientific, Cat # 3431D76) on ice. The homogenate was loaded into a 30 mL thick polycarbonate ultracentrifuge tube (Beckman Coulter, Cat # 355631). 9 mL of sucrose solution (1.8 M sucrose, 3 mM MgAc^2^, 1 mM DTT, 10 mM Tris-HCl in DEPC-treated water) was added to the bottom of the tube with the homogenate and centrifuged at 107,000 g for 2.5 hours at 4°C. Supernatant was aspirated, and the nuclei containing pellet was incubated in 250 uL of DEPC-treated water-based PBS for 20 min on ice before resuspending the pellet. The nuclear suspension was filtered twice through a 30 um cell strainer. Nuclei were counted using a hemocytometer and diluted to 2,000 nuclei/uL before performing single-nuclei capture on the 10X Genomics Single-Cell 3’ system. Usually, the target capture of 3,000 nuclei per sample was used; the 10x capture and library preparation protocol was used without modification. Single-nucleus libraries from individual samples were pooled and sequenced on the NovaSeq 6000 machine (average depth 60,000 reads/nucleus).

#### snRNA-seq data processing with 10X Genomics CellRanger software and data filtering

For library demultiplexing, fastq file generation and read alignment and UMI quantification, CellRanger software v 1.3.1 was used. CellRanger was used with default parameters, except for using pre-mRNA reference file (ENSEMBL GRCh38) to insure capturing intronic reads originating from pre-mRNA transcripts abundant in the nuclear fraction.

Individual expression matrices containing numbers of Unique molecular identifiers (UMIs) per nucleus per gene were filtered to retain nuclei with at least 400 genes expressed and less than 10% of total UMIs originating from mitochondrial and ribosomal RNAs. Individual matrices were combined prior to pre-processing and clustering with Seurat.

#### snRNA-seq dataset integration, dimensionality reduction, UMAP embedding, clustering and cell type identification

All of the following bioinformatics analysis steps are documented in an R script available at https://doi.org/10.5281/zenodo.7245297. In order to integrate snRNA-seq datasets, we utilized Harmony (*31*) integration using the 10x Genomics chemistry version as the grouping variable. Downstream data preprocessing, normalization, variable feature selection and PCA was performed using the standard Seurat pipeline (*32*). Selection of significant principal components was done using the elbow method. The selected components were used to perform UMAP embedding and clustering using the Louvain method. The identity of specific lineages and cell types was determined based on expression of known marker genes, as is shown in Figure 1 and Figure S1.

### Sex determination

To determine the sex of individuals for which sex information was not available, we aggregated gene expression of all nuclei by individual and plotted individual-wise expression of the following genes: *XIST*, *DDX3Y*, *KDM5D*, *USP9Y*, *ZFY*, *EIF1AY*, *UTY*.

#### Trajectory reconstruction and isolation of individual lineages

Seurat UMAP coordinates were imported into monocle3 (*33*) for trajectory reconstruction. learn_graph function with custom graph_control options was used to construct the trajectory graph. We noticed that while the original trajectory graph generated by monocle3 corresponded to the major cell lineages, it failed to connect some nodes that passed through populations of cells expressing shared lineage markers. Moreover, some trajectory branches did not correspond to biologically interpretable lineage progression, specifically the branches connecting two mature neuronal cell types containing only adult cells. We corrected these issues by modifying the trajectory according to the following principles: 1) if two terminal nodes failed to be connected but were passing through populations of cells expressing known lineage-specific markers (such as *RORB* for layer 4, *TLE4*/*SEMA4A* for layer 6b, *CUX2* for layer 2-3 and *CUX1* for layer 5-6-IT), we connected these nodes 2) if a branch connected nodes located in two mature cell types, we omitted this branch and 3) based on the first two principles, we isolated the shortest path between the node in the neural progenitor/radial glia cluster and the node in the mature cell type cluster.

#### Identification of lineage-specific dynamically expressed genes

First, we selected trajectory branches corresponding to specific lineages, as well as the cells along the branches. For the interneuron trajectory analysis, we only selected MGE or CGE cells from the GE progenitors cluster to analyze MGE and CGE-derived INs, respectively. Then, monocle3’s Moran’s test (graph_test function) was used to identify genes that are dynamically expressed in each lineage. We modified graph_test function to utilize Moran’s test with covariates to ensure that our results are not affected by uneven contribution of cells from male and female subjects, different brain regions, as well as cells postmortem interval and 10x chemistry. We selected genes with adjusted p value < 0.05 as statistically significant dynamically expressed genes. To identify lineage-specific genes, we first compressed the single-cell expression data along each lineage by using a sliding window along pseudotime and averaging expression of neighboring cells for each gene. We generated 500 meta-cells in each lineage using this approach. Then, we fit the expression of each gene using a generalized linear model and the following formula: expression ∼ splines::ns(pseudotime, df=3). Then, we calculated the area under the curve for the smoothed expression/pseudotime plot for each gene in each lineage across intervals of the sliding window. The difference of under the curve between the lineage of interest and all other lineages was used to rank genes according to their lineage specificity. Moran’s p value < 0.05 and an expression difference of at least 20% in one section of the sliding window was used to define lineage-specific genes.

#### Analysis of single-cell ATAC-seq data and snRNA-seq/scATAC-seq integration

Four scATAC-seq datasets were first remapped to the same hg38 genome reference. Then, a minimal non-overlapping consensus peak set was created based on the peaks from all datasets, and ATAC-seq counts were mapped on this set of peaks using Signac (*34*), and the datasets were combined. Then, gene activity matrix for the combined dataset was generated by counting ATAC peaks in the promoter region and the gene body, using the same parameters as used by the Signac package. For mapping scATAC-seq data on the snRNA-seq dataset, we first integrated the two modalities using Seurat’s FindTransferAnchors and the canonical correlation analysis (cca). We used the expression and gene activity of genes variable in the snRNA-seq datasets to perform cca and then used the TransferData function to map the scATAC-seq data on the snRNA-seq space followed by Harmony processing to regress the effect of different scATAC-seq and snRNA-seq chemistries. To map scATAC-seq profiles to the UMAP space and clusters we generated using snRNA-seq data, we identified 100 nearest neighbors for each scATAC-seq cell in the combined snRNA-seq/scATAC-seq space and then calculated the UMAP coordinates and cluster membership in the snRNA-seq space. To validate the accuracy of this procedure, we checked for the specificity of gene activity of cell type markers, as well as for age distribution. This integration and mapping procedure was repeated for the three major lineage classes (excitatory neurons, interneurons and macroglial cells).

#### SCENIC+ analysis

SCENIC+ requires single-cell transcriptomic and scATAC-seq data mapped to the same category (e.g. cluster) and also recommends generating pseudobulk scATAC-seq profiles prior to the analysis. In order to prepare our data for SCENIC+ analysis, we first selected ATAC-seq cells along the lineage trajectories using a sliding window approach and keeping the cells in cell type-specific clusters. Then, we generated 2500 meta-cell pseudobulk ATAC-seq profiles using the sliding window along each trajectory and summing all ATAC counts. We also generated 2500 meta-cells for the corresponding lineage-specific snRNA-seq profiles and restricted the analysis to lineage and branch-specific genes relevant to each lineage. In order to generate pseudo-multiome profiles from separate snRNA-seq and scATAC-seq datasets, we sorted cells into 10 bins based on the pseudotime progression. These pseudotime bins were also used to identify differentially accessible regions of chromatin and cis-regulatory topics using cisTopic (*35*), which was used with default settings, except for setting the differential features threshold to 25%. After generating pseudo-multiome profiles, we performed SCENIC+ analysis as described in the tutorial. Significant enhancer-transcription factor-gene relationships in each lineage were exported as the final result.

#### Identification of sex and region-enriched dynamically expressed genes

To identify male and female-enriched genes in each lineage, we selected cells from only males or females within each lineage and first performed Moran’s I test separately for male and female data. Then, we compressed the data and calculated area under the curve for male and female gene expression. Genes with Moran’s I statistic >= 0.1, adjusted Moran’s p value<0.05 and the area under curve difference between male and female expression >= 50 were considered sex-specific in each given lineage.

#### Gene ontology analysis

We used ShinyGO (*36*) to perform gene ontology analysis using genes expressed in each lineage as the background gene list. In order to reduce redundancy of the identified GO terms, all significant (adjusted p value < 0.05) terms were used as input to Revigo (*37*) in case more than 10 pathways were identified. The value of the resulting gene list of 0.4 was used. The -log10(p value) and fold enrichment for the resulting non-redundant GO processes were reported.

#### Analysis of enrichment of disease risk genes

We intersected disease risk gene lists with our list of lineage-specific genes, as well as genes enriched in male and female developmental lineages. We calculated hypergeometric p values for each overlap, using genes expressed in each lineage as the background.

### Data visualization

Cell type, gene expression and lineage trajectories for each lineage can be visualized at https://pre-postnatal-cortex.cells.ucsc.edu.

#### MERSCOPE spatial transcriptomics

Sample preparation was performed according to manufacturer’s instructions (MERSCOPE Fresh and Fixed Frozen Tissue Sample Preparation User Guide, Doc. number 91600002). Briefly, fresh snap frozen tissue with a high RNA integrity number (RIN>8) were sectioned (10um thick) using a cryostat and mounted on MERSCOPE functional slides. Sections where then fixed and stored at 70% ethanol for up to two weeks. Sections went through autofluorescence quenching under UV light for 3 hours using the MERSCOPE Photo-bleacher instrument. A Pre-designed panel mix (140 genes) focused on early emerging excitatory lineage-specific genes based on the single-nuclei analysis were used for probe hybridization. Hybridizations were performed at 37°C for up to 48 hours in a humid environment. Post prob hybridization, sections were fixed using formamide and embedded in gel. After gel embedding, tissue samples were cleared using a clearing mix solution supplemented with proteinase K for 24-48 hours at 37°C until no visible tissue was evident in the gel. After clearing was completed, sections were stained for DAPI and PolyT and fixed with formamide prior to imaging. No additional cell boundary stainings were used. The MERSOPE imaging process was done according to the MERSCOPE Instrument Site Preparation Guide (Doc. Number 91500001). Briefly, an imaging kit was thawed at 37°C for 45 minutes, activated and loaded into the MERSCOPE instrument. The flow chamber was then assembled, fluidics were primed, flow chamber filled with liquid and a low-resolution image was taken. Based on DAPI staining, an ROI was chosen for the full imaging experiment. After imaging was complete, data was processed using MERSCOPE proprietary software. Further analysis, visualization, and integration of spatial data, was done using Seurat v5 (Source: vignettes/spatial_vignette_2.Rmd). Putative neuronal layer localization was predicted from co-localization with referenced markers at relevant developmental stages.

---

**Table S1. Sample and nuclei metadata.**

**Table S2. Lineage and branch-specific genes.**

**Table S3. Results of eGRN analysis using SCENIC+.**

**Table S4. Results of region-specific gene expression analysis.**

**Table S5. Sex-enriched developmentally regulated genes.**

**Table S6. Lineage- and sex-specific disease risk genes.**

**Figure S1.**
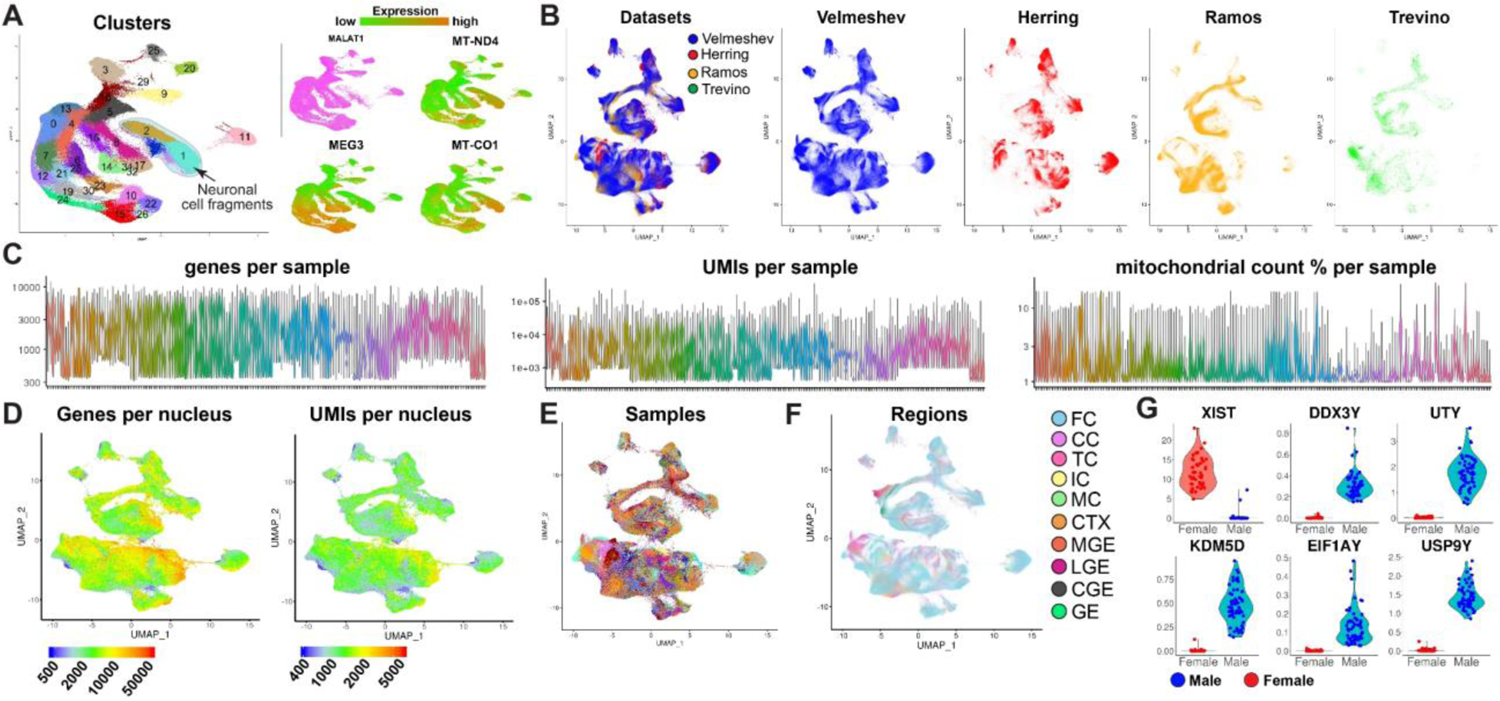
Technical and biological characteristics of the combined snRNA-seq dataset. **A)** Identification of the clusters containing neuronal debris. **B)** Integration of the current dataset with previously published datasets. **C)** Gene and UMI counts per nucleus, as well as mitochondrial reads ratio across all samples. **D)** Gene and UMI counts per nucleus across all cell types. **E-F)** Distribution of nuclei from different samples and regions. FC-frontal/prefrontal cortex, CC-cingulate cortex, TC-temporal cortex, IC-insular cortex, MC-motor cortex, CTX-cortex. **G)** Expression of sex-specific genes used to determine sex of samples with unknown status.

**Figure S2.**
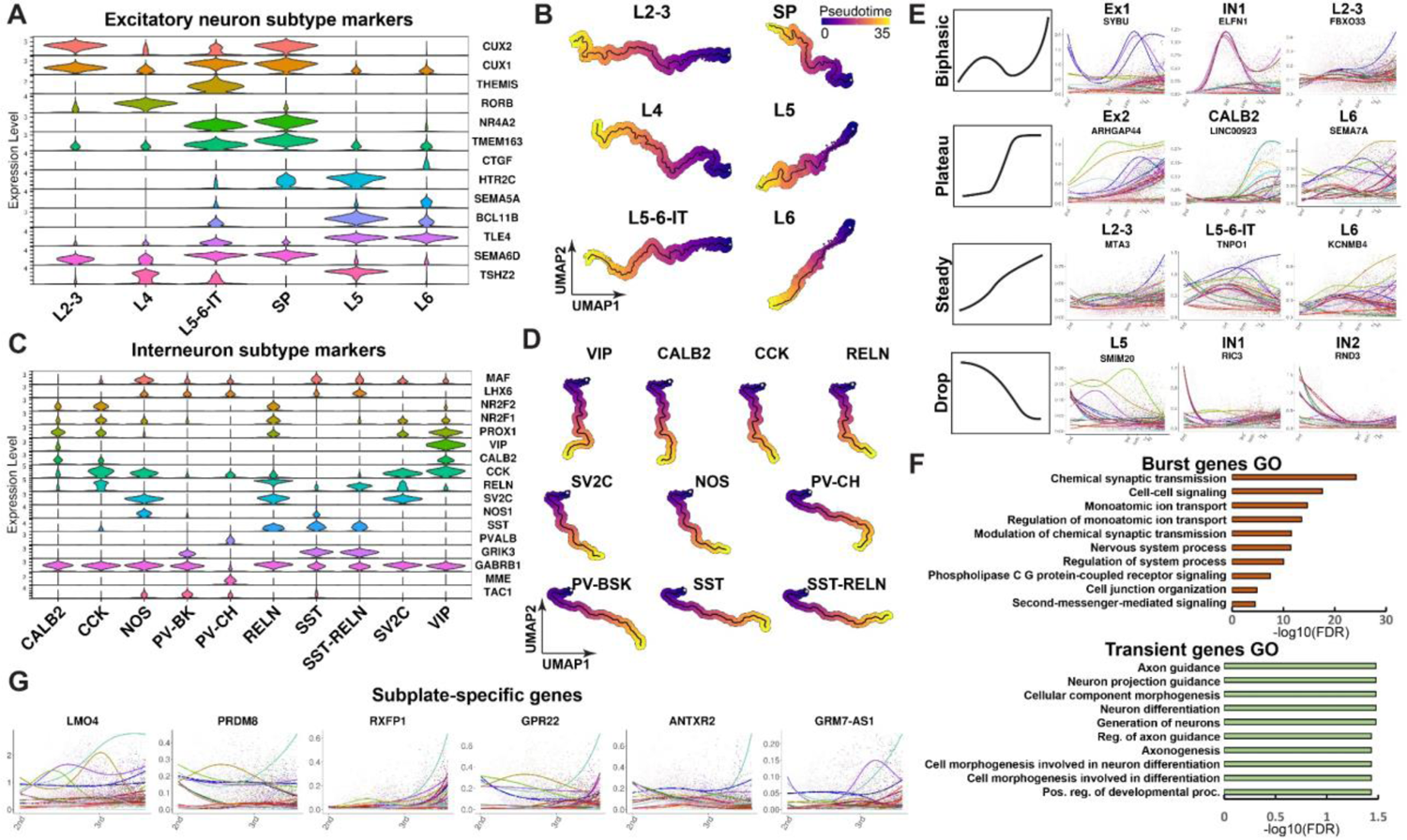
Excitatory neuron and interneuron lineage analysis. **A)** Expression of cortical excitatory neuron marker genes used to determine excitatory neuron lineages. **B)** Isolated lineages trajectories for excitatory neuron subtypes. **C)** Markers of interneuron subtypes. **D)** Isolated interneuron trajectories. **E)** Examples of biphasic, plateau, steady and drop expression of lineage and branch-specific genes. **F)** GO pathways enriched for burst and transient neuronal genes. **G)** Top subplate-specific dynamically expressed genes.

**Figure S3.**
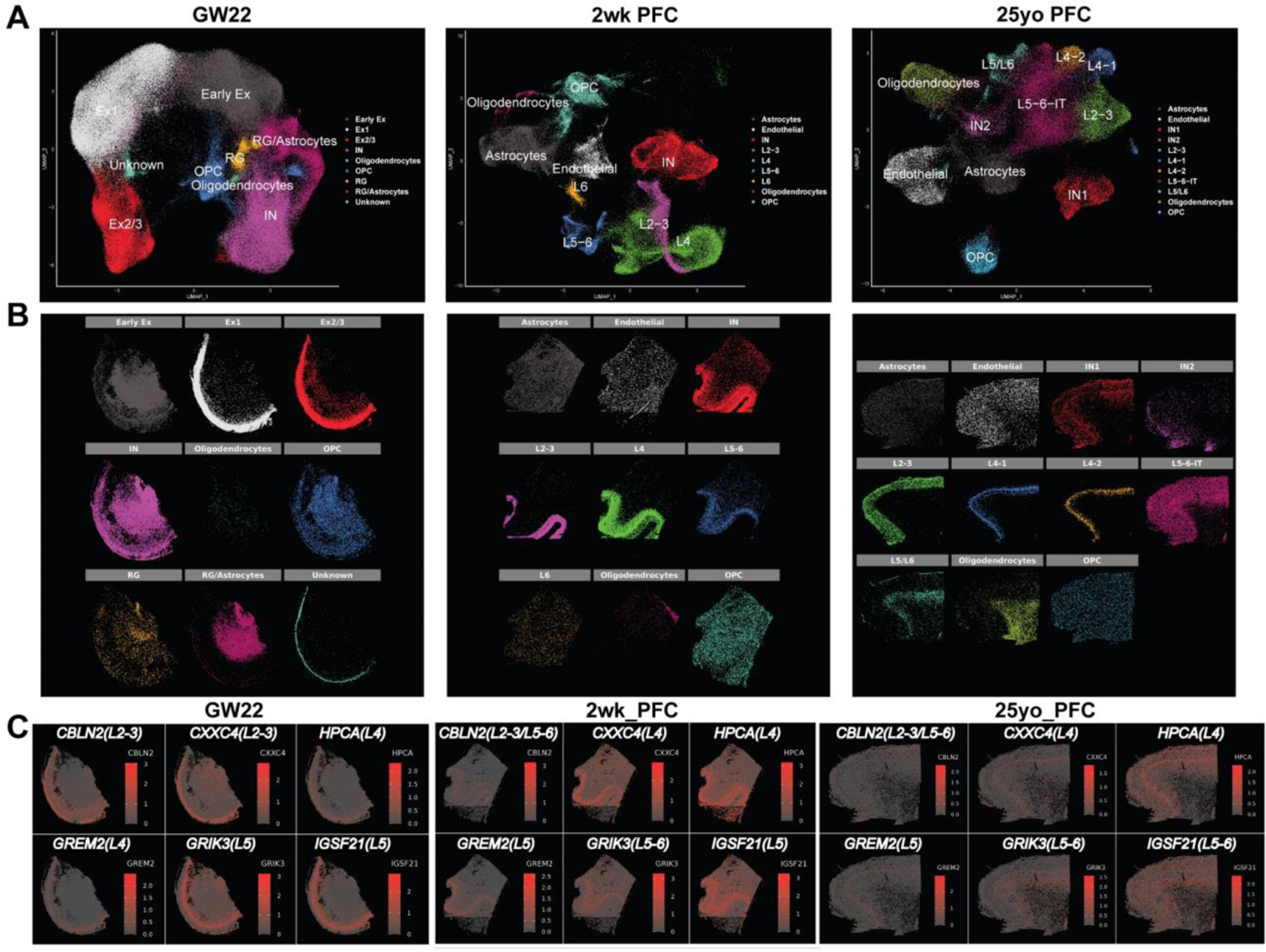
Spatial transcriptomic analysis of lineage-specific genes across development. **A)** UMAP embedding of annotated clusters. **B)** Spatial localization patterns of individual clusters (cluster colors and spatial location correspond with Fig. 2g). **C)** Spatiotemporal expression of layer-specific markers.

**Figure S4.**
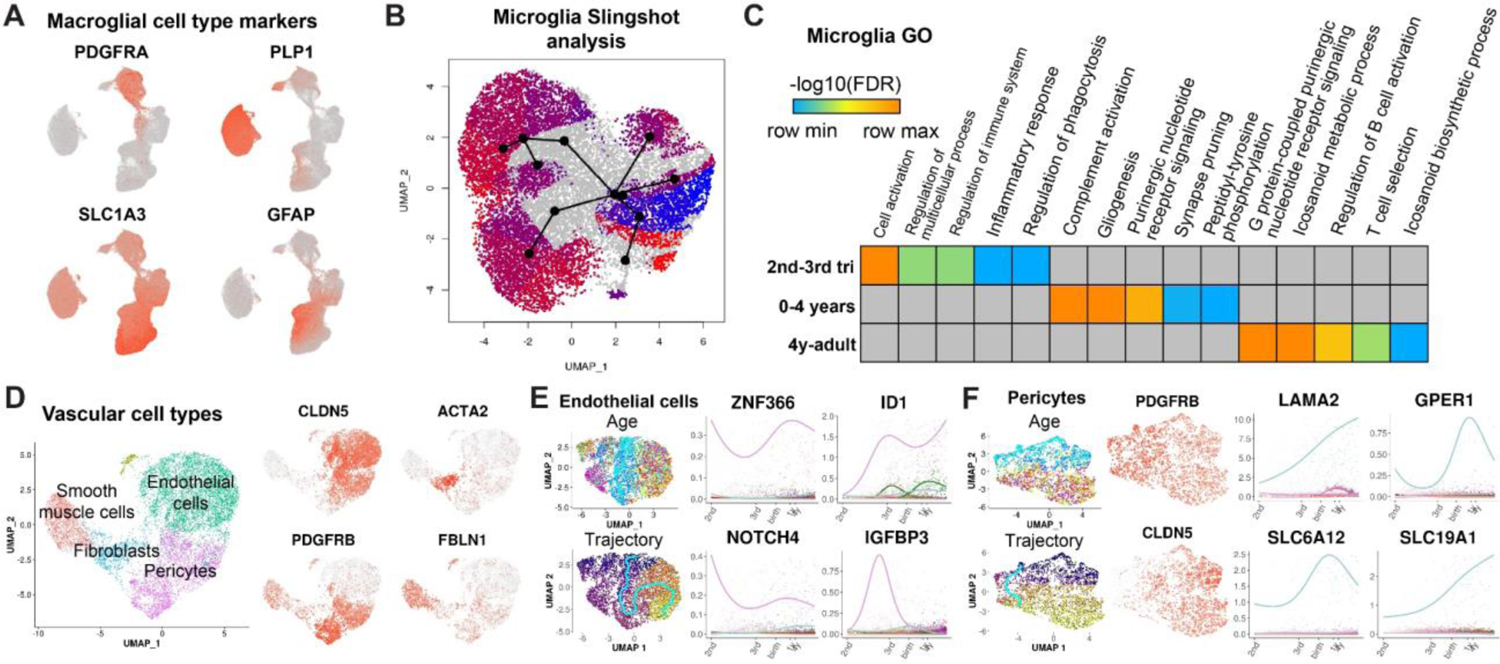
Analysis of glial and vascular lineages. **A)** Markers of OPCs, oligodendrocytes, fibrous and protoplasmic astrocytes **B)** Slingshot analysis of microglial lineage trajectories. **C)** Gene ontology analysis developmental microglia genes. **D)** Analysis of vascular cell types. **E-F)** Trajectory analysis of endothelial cells and pericytes.

**Figure S5.**
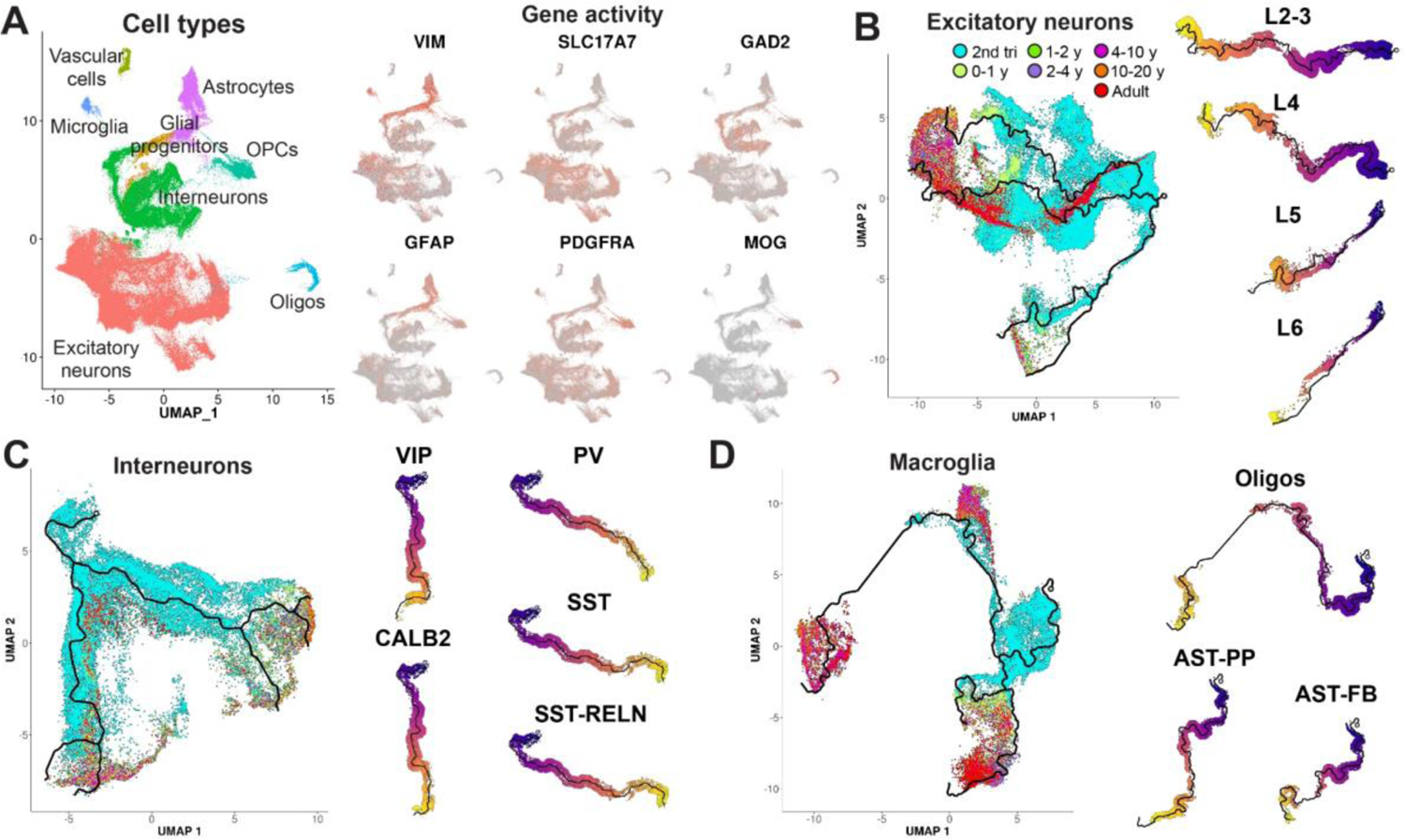
Mapping developmental scATAC-seq to specific lineage trajectories. **A)** Gene activities of cell type-specific marker genes. **B-D)** Age distribution and selection of ATAC-seq cells for specific lineages of excitatory neurons **(B)**, interneurons **(C)** and macroglial cells **(D)**.

**Figure S6.**
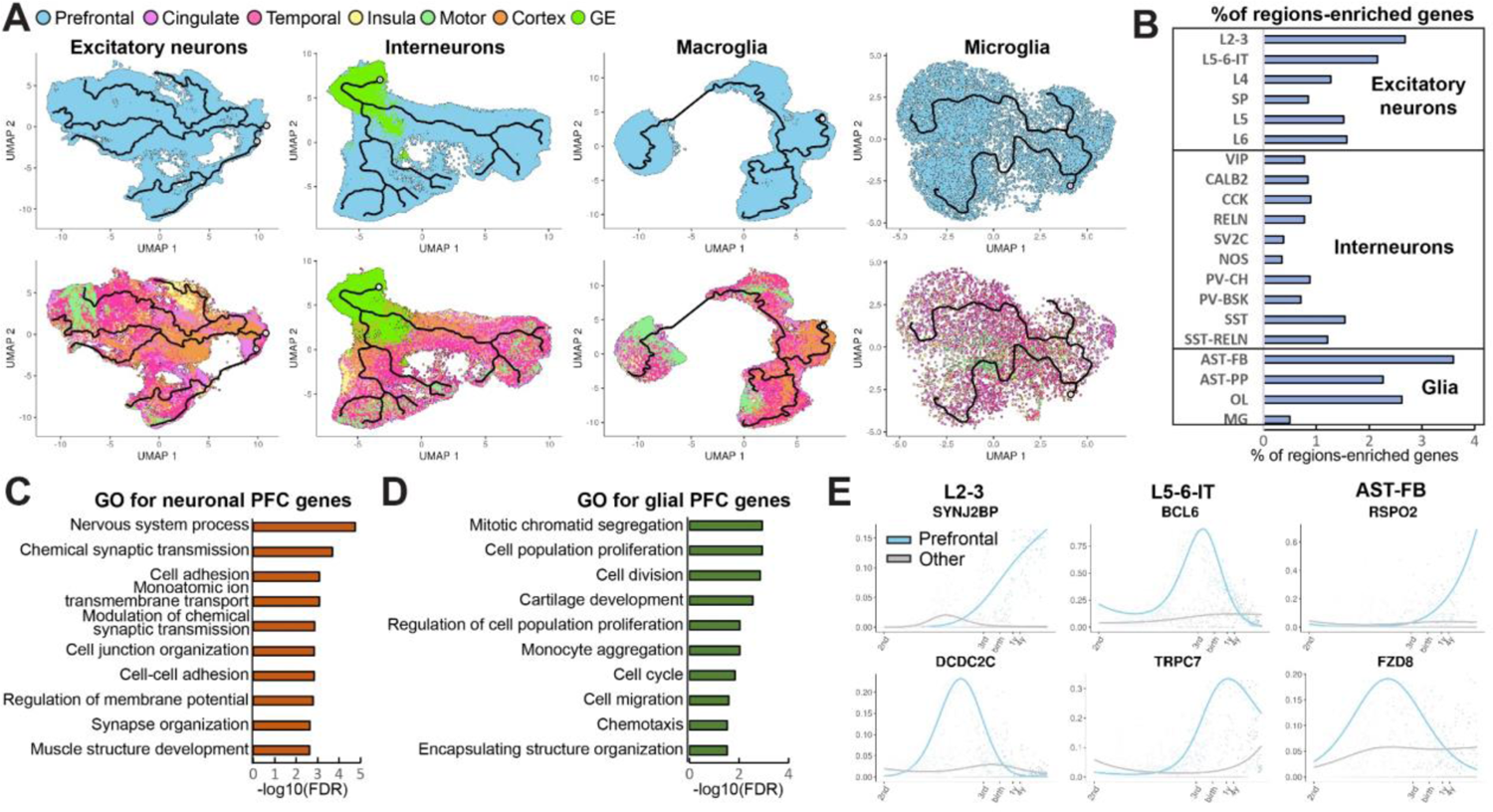
Frontal cortex-specific developmental programs. **A)** Cells from the frontal/prefrontal cortex and other cortical regions in the excitatory neuron, interneuron, macroglial and microglial lineages. **B)** Number of PFC-specific genes in neuronal and glial lineages relative to the total number of genes expressed in each lineage. **C-D)** Gene ontology analysis of PFC-specific genes in neuronal and glial lineages. **E)** Examples of top genes enriched in the PFC in specific lineages.

**Figure S7.**
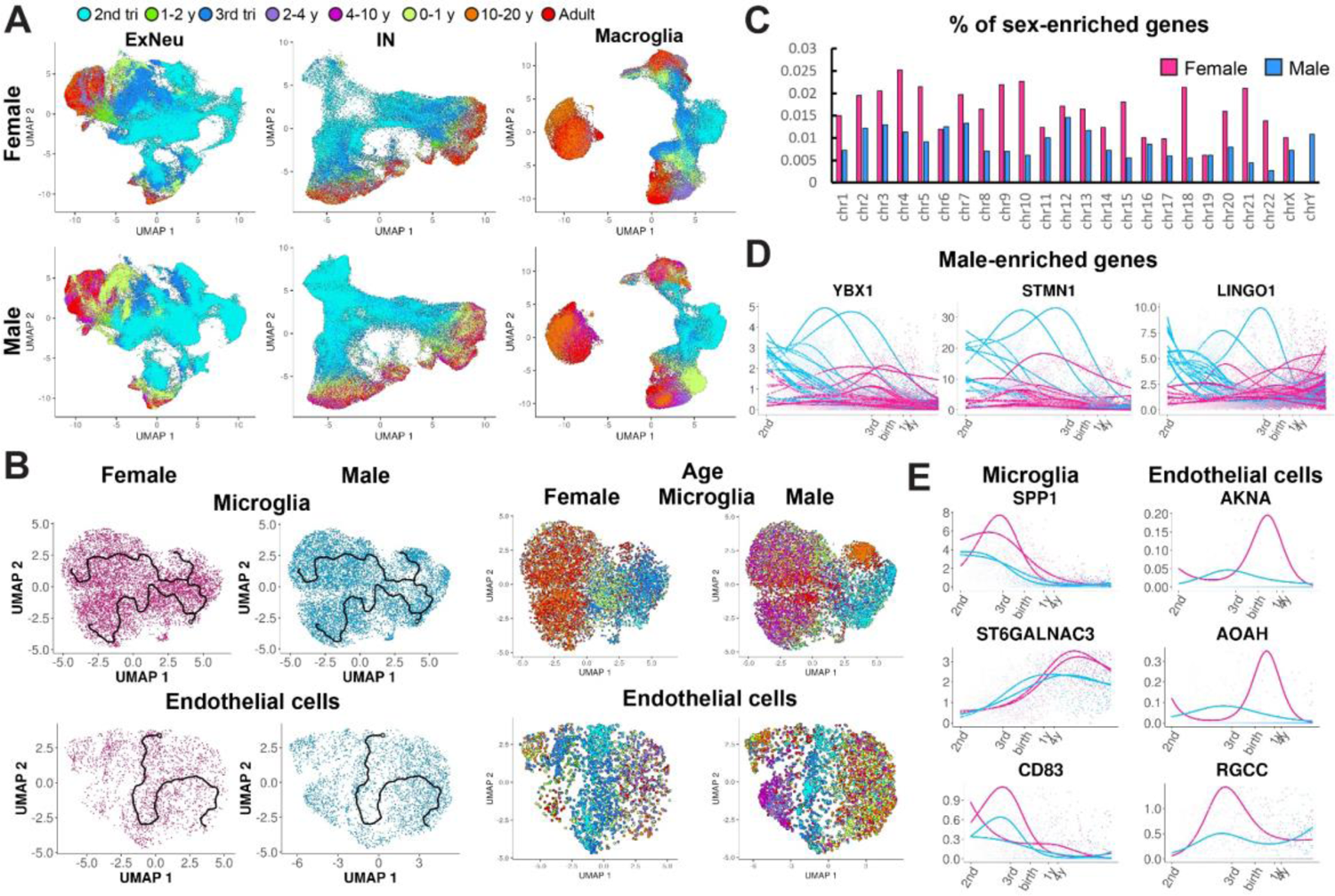
Analysis of sex and region-enriched genes during microglia and endothelial cell development. **A)** Female and male microglia and endothelial cell trajectories. **B)** relative number of sex-specific genes per chromosome. **C)** Examples of top male-enriched genes. **D)** Female and male trajectories in microglia and endothelial cells. **E)** Top female-enriched genes expressed in microglia and endothelial cells.

