## Supplementary material for "Single-cell analysis of prenatal and postnatal human cortical development"

**One Sentence Summary:** Single-cell transcriptomic atlas of human cortical development identifies lineage and sex-specific programs and their implication in brain disorders.

**Authors:** Dmitry Velmeshev*#[1,2,3], Yonatan Perez*[1,2], Zihan Yan[3], Jonathan E. Valencia[4], David R. Castaneda-Castellanos[5], Li Wang[1,2], Lucas Schirmer[6,7,8], Simone Mayer[1,2,9], Brittney Wick[10], Shaohui Wang[1,2], Tomasz Jan Nowakowski[11], Mercedes Paredes[2], Eric J Huang[1,12], Arnold R Kriegstein[1,2#]

**Affiliations:** *[1]Eli and Edythe Broad Center of Regeneration Medicine and Stem Cell Research, University of California, San Francisco, CA 94143; [2]Department of Neurology, University of California, San Francisco, CA 94143, [3]Department of Neurobiology, Duke University School of Medicine, Durham, NC 27710; [4]Curio Bioscience, 4030 Fabian Way, Palo Alto, CA 94303, [5]Vizgen Inc. 61 Moulton Street Cambridge, MA 02138, [6]Division of Neuroimmunology, Department of Neurology, Medical Faculty Mannheim, Heidelberg University, Mannheim, Germany, 68167; [7]Mannheim Center for Translational Neuroscience and Institute for Innate Immunoscience, Medical Faculty Mannheim, Heidelberg University, Mannheim, Germany, 68167; [8]Interdisciplinary Center for Neurosciences, Heidelberg University, Heidelberg, Germany, 68167; [9]Hertie Institute for Clinical Brain Research, University of Tübingen, Tübingen, Germany 72076; [10]UC Santa Cruz Genomics Institute, Santa Cruz, CA 95060, [11]Department of Neurological Surgery, University of California, San Francisco, CA 94143; [12]Department of Pathology, University of California, San Francisco, CA 94115.*

**Table S1. Sample and nuclei metadata.**

**Table S2. Lineage and branch-specific genes.**

**Table S3. Results of eGRN analysis using SCENIC+.**

**Table S4. Results of region-specific gene expression analysis.**

**Table S5. Sex-enriched developmentally regulated genes.**

**Table S6. Lineage- and sex-specific disease risk genes.**

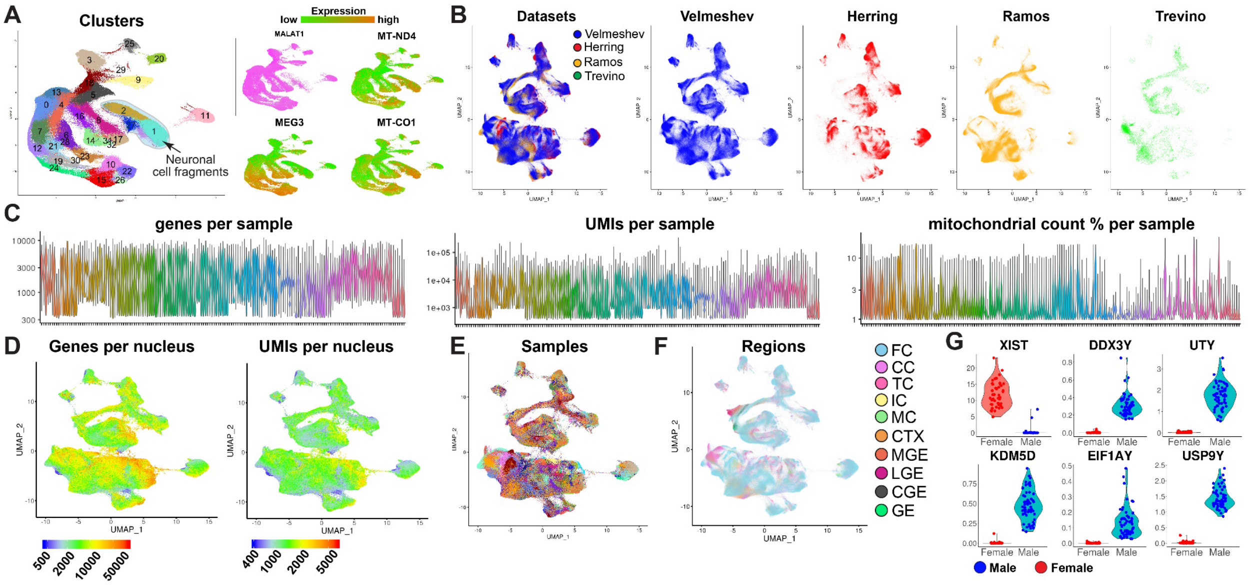


**Figure S1. Technical and biological characteristics of the combined snRNA-seq dataset. A)** Identification of the clusters containing neuronal debris. **B)** Integration of the current dataset with previously published datasets. **C)** Gene and UMI counts per nucleus, as well as mitochondrial reads ratio across all samples. **D)** Gene and UMI counts per nucleus across all cell types. **E-F)** Distribution of nuclei from different samples and regions. FC-frontal/prefrontal cortex, CC-cingulate cortex, TC-temporal cortex, IC-insular cortex, MC-motor cortex, CTX-cortex. **G)** Expression of sex-specific genes used to determine sex of samples with unknown status.

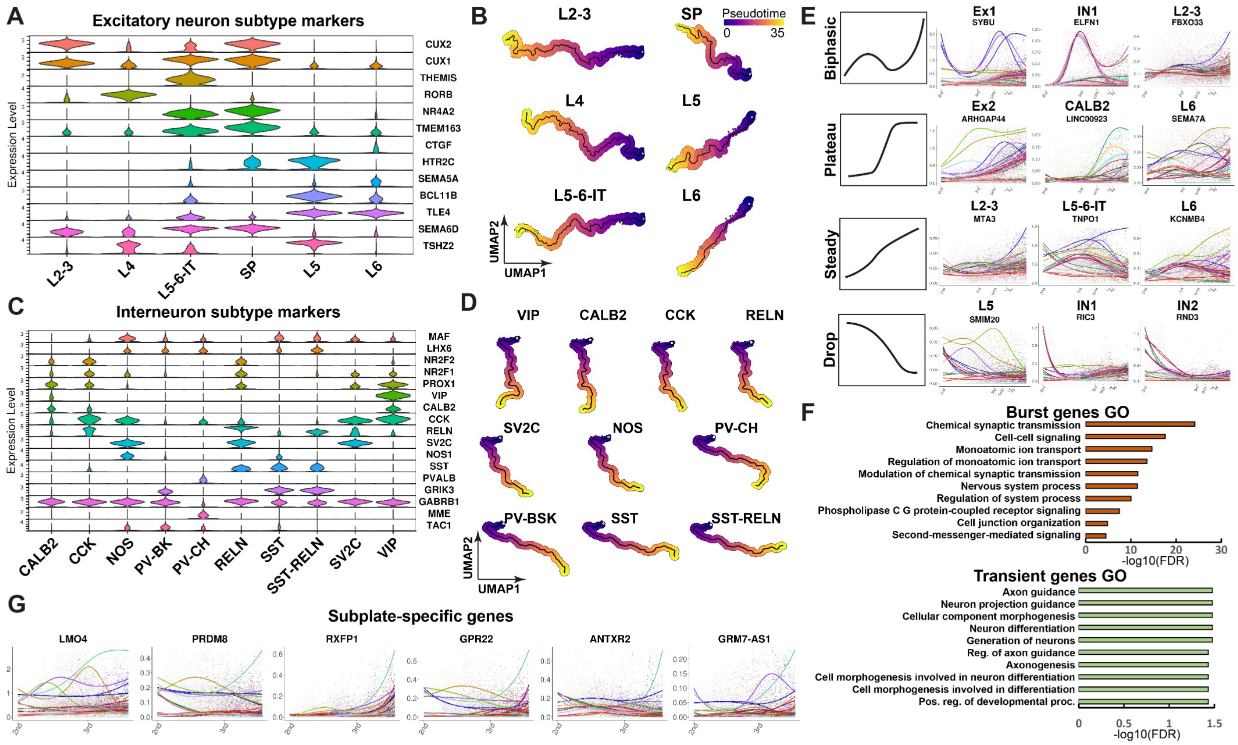


**Figure S2. Excitatory neuron and interneuron lineage analysis. A)** Expression of cortical excitatory neuron marker genes used to determine excitatory neuron lineages. **B)** Isolated lineages trajectories for excitatory neuron subtypes. **C)** Markers of interneuron subtypes. **D)** Isolated interneuron trajectories. **E)** Examples of biphasic, plateau, steady and drop expression of lineage and branch-specific genes. **F)** GO pathways enriched for burst and transient neuronal genes. **G)** Top subplate-specific dynamically expressed genes.

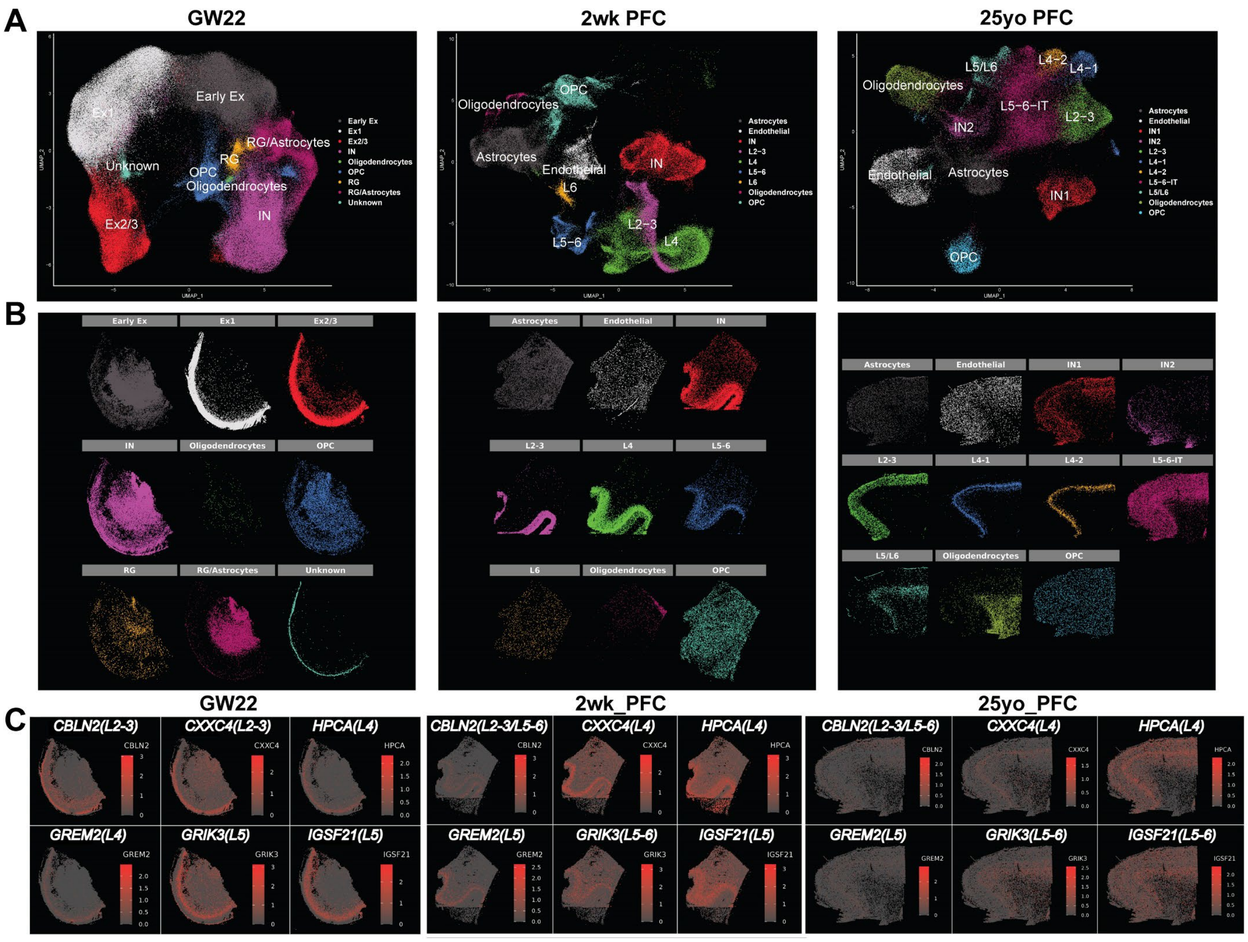


**Figure S3. Spatial transcriptomic analysis of lineage-specific genes across development. A)** UMAP embedding of annotated clusters. **B)** Spatial localization patterns of individual clusters (cluster colors and spatial location correspond with Fig. 2g). **C)** Spatiotemporal expression of layer-specific markers.

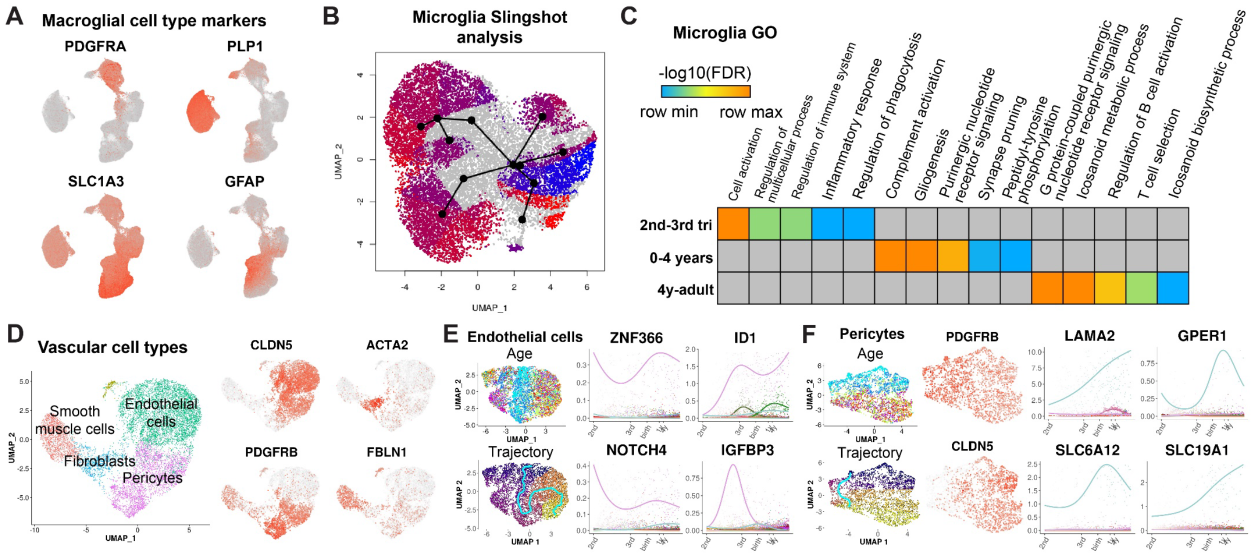


**Figure S4. Analysis of glial and vascular lineages. A)** Markers of OPCs, oligodendrocytes, fibrous and protoplasmic astrocytes **B)** Slingshot analysis of microglial lineage trajectories. **C)** Gene ontology analysis developmental microglia genes. **D)** Analysis of vascular cell types. **E-F)** Trajectory analysis of endothelial cells and pericytes.

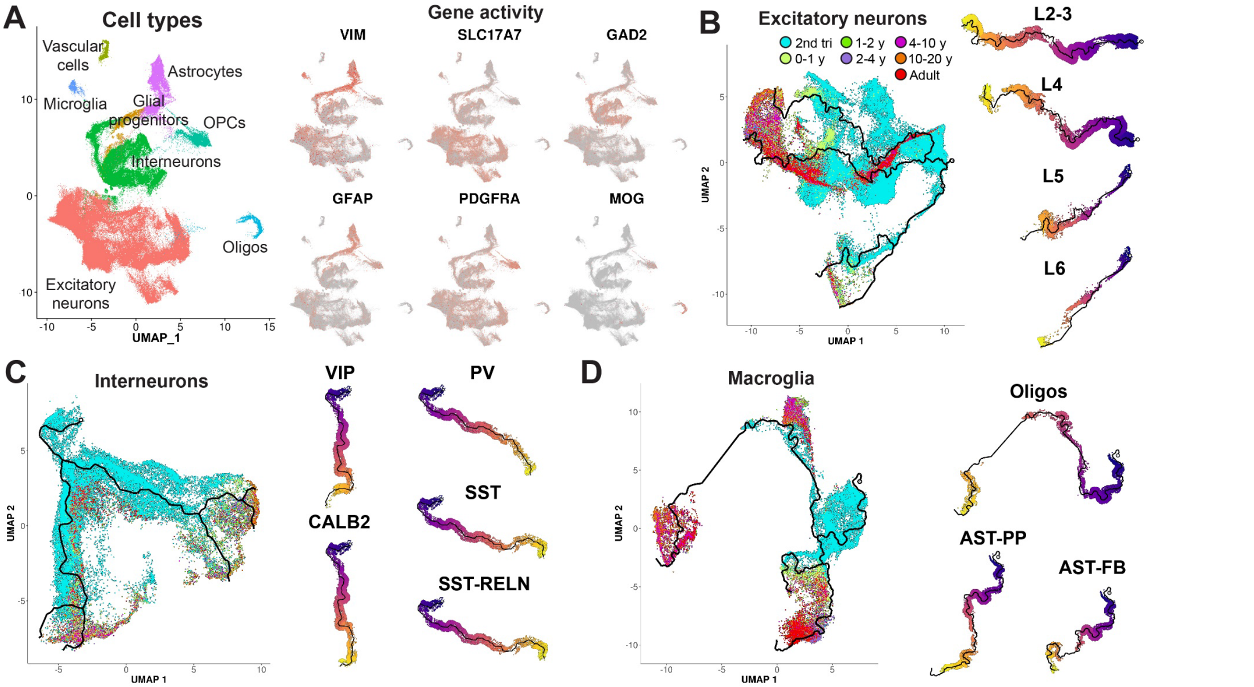


**Figure S5. Mapping developmental scATAC-seq to specific lineage trajectories. A)** Gene activities of cell type-specific marker genes. **B-D)** Age distribution and selection of ATAC-seq cells for specific lineages of excitatory neurons **(B)**, interneurons **(C)** and macroglial cells **(D)**.

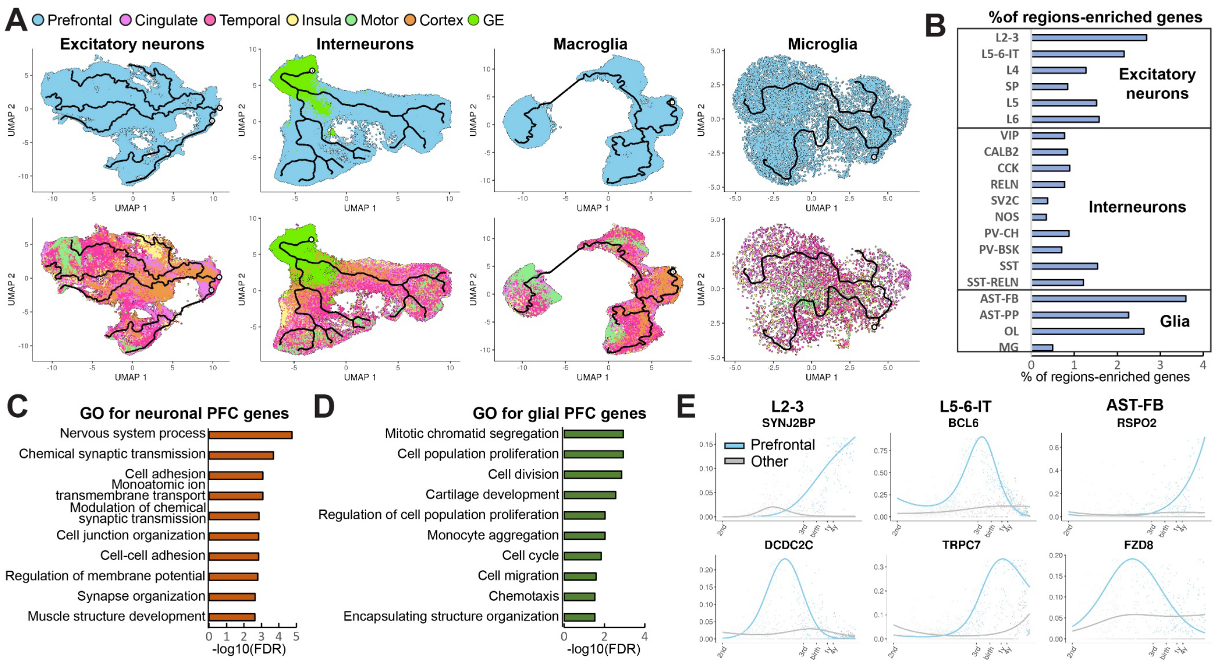


**Figure S6. Frontal cortex-specific developmental programs. A)** Cells from the frontal/prefrontal cortex and other cortical regions in the excitatory neuron, interneuron, macroglial and microglial lineages. **B)** Number of PFC-specific genes in neuronal and glial lineages relative to the total number of genes expressed in each lineage. **C-D)** Gene ontology analysis of PFC-specific genes in neuronal and glial lineages. **E)** Examples of top genes enriched in the PFC in specific lineages.

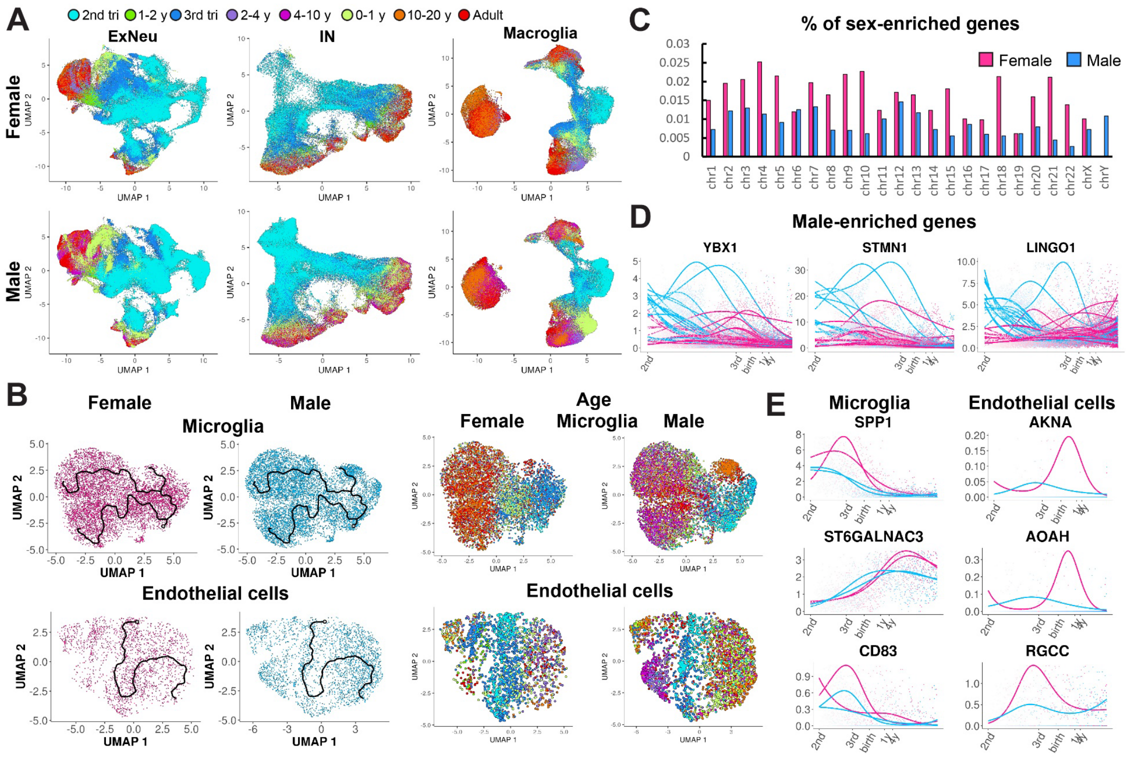


**Figure S7. Analysis of sex and region-enriched genes during microglia and endothelial cell development. A)** Female and male microglia and endothelial cell trajectories. **B)** relative number of sex-specific genes per chromosome. **C)** Examples of top male-enriched genes. **D)** Female and male trajectories in microglia and endothelial cells. **E)** Top female-enriched genes expressed in microglia and endothelial cells.
